# Global analysis of adenylate-forming enzymes reveals β-lactone biosynthesis pathway in pathogenic *Nocardia*

**DOI:** 10.1101/856955

**Authors:** Serina L. Robinson, Barbara R. Terlouw, Megan D. Smith, Sacha J. Pidot, Tim P. Stinear, Marnix H. Medema, Lawrence P. Wackett

## Abstract

Enzymes that cleave ATP to activate carboxylic acids play essential roles in primary and secondary metabolism in all domains of life. Class I adenylate-forming enzymes share a conserved structural fold but act on a wide range of substrates to catalyze reactions involved in bioluminescence, nonribosomal peptide biosynthesis, fatty acid activation, and β-lactone formation. Despite their metabolic importance, the substrates and catalytic functions of the vast majority of adenylate-forming enzymes are unknown without tools available to accurately predict them. Given the crucial roles of adenylate-forming enzymes in biosynthesis, this also severely limits our ability to predict natural product structures from biosynthetic gene clusters. Here we used machine learning to predict adenylate-forming enzyme function and substrate specificity from protein sequence. We built a web-based predictive tool and used it to comprehensively map the biochemical diversity of adenylate-forming enzymes across >50,000 candidate biosynthetic gene clusters in bacterial, fungal, and plant genomes. Ancestral enzyme reconstruction and sequence similarity networking revealed a ‘hub’ topology suggesting radial divergence of the adenylate-forming superfamily from a core enzyme scaffold most related to contemporary aryl-CoA ligases. Our classifier also predicted β-lactone synthetases in novel biosynthetic gene clusters conserved across >90 different strains of *Nocardia*. To test our computational predictions, we purified a candidate β-lactone synthetase from *Nocardia brasiliensis* and reconstituted the biosynthetic pathway *in vitro* to link the gene cluster to the β-lactone natural product, nocardiolactone. We anticipate our machine learning approach will aid in functional classification of enzymes and advance natural product discovery.

## INTRODUCTION

Adenylation is a widespread and essential reaction in nature to transform inert carboxylic acid groups into high energy acyl-AMP intermediates. Class I adenylate-forming enzymes catalyze reactions for natural product biosynthesis, firefly bioluminescence, and the activation of fatty acids with coenzyme A (1). The conversion of acetate to acetyl-CoA by a partially purified acetyl-CoA ligase was first described by Lipmann in 1944 (2). Since then, enzymes with this conserved structural fold have been found to activate over 200 different substrates including aromatic, aliphatic, and amino acids. To encompass the major functional enzyme classes, the term ‘ANL’ superfamily was proposed based on the <u>A</u>cyl-CoA ligases, <u>N</u>onribosomal peptide synthetases (NRPS), and <u>L</u>uciferases (3).

Most ANL superfamily enzymes catalyze two-step reactions: adenylation followed by thioesterification. During the thioesterification step, ANL enzymes undergo a dramatic conformational change involving a 140° domain rotation of the C-terminal domain (3). Thioester bond formation results from nucleophilic attack, typically by a phosphopantetheine thiol group. A notable exception to phosphopantetheine is the use of molecular oxygen by firefly luciferase to convert D-luciferin to a light-emitting oxidized intermediate (3). Other interesting exceptions include functionally-divergent ANL enzymes within the same pathway that catalyze the adenylation and thioesterification partial reactions separately (4, 5). The ANL superfamily has recently expanded to include several new classes of enzymes including the fatty-acyl AMP ligases (6), aryl polyene adenylation enzymes (7), and β-lactone synthetases (8). Strikingly, an ANL enzyme in cremeomycin biosynthesis was shown to use nitrite to catalyze late stage N-N bond formation in diazo-containing natural products (9). The discovery of such novel enzymatic reactions nearly 75 years after the Lipmann’s initial report suggests the ANL superfamily still has unexplored catalytic potential, particularly in specialized metabolism.

No computational tools currently exist for prediction and functional classification of ANL enzymes at the superfamily level. Still, the development of one previous platform, which is no longer supported or available, showed that this class of enzymes is amenable to computational predictions of substrate and function (10). Previously, bioinformatics tools were also developed to predict substrates for NRPS adenylation (A) domains (11, 12). Genome mining approaches using NRPS A domain prediction tools have proved useful to access the biosynthetic potential of unculturable organisms and link ‘orphan’ natural products with their biosynthetic gene clusters. For example, NRPS A domain predictions guided the discovery of the biosynthetic machinery for the leinamycin family of natural products (13) and enabled sequence-based structure prediction of several novel lipopeptides by Zhao et al. (14). However, Zhao et al. reported limitations in existing tools in that they could not predict the chain length of lipid tails incorporated into lipopeptides. Lipid tails are prevalent in natural products and are incorporated by fatty acyl-AMP or acyl-CoA ligase enzymes, both in the ANL superfamily. These enzymes are among the most well-studied subclasses of ANL enzymes. For less-studied subclasses, enzyme functions and substrates are even more challenging to predict. Hence, a computational tool encompassing all classes of ANL enzymes would constitute a major step towards more accurate structural prediction of natural product scaffolds.

Here, we used machine learning to develop a predictive platform for ANL superfamily enzymes and map their substrate-and-function landscape across 50,064 candidate biosynthetic gene clusters in bacterial, fungal, and plant genomes. We detected candidate β-lactone synthetases in uncharacterized biosynthetic gene clusters from pathogenic *Nocardia* spp. and experimentally validated the gene cluster *in vitro* to link it to the orphan β-lactone compound, nocardiolactone. Overall, this research provides a proof-of-principle towards the use of machine learning for classification of enzyme substrates to guide natural product discovery.

## RESULTS

### Machine learning can accurately predict ANL enzyme function and substrate specificity

A global analysis of protein family domains revealed that ANL superfamily enzymes (PF00501; AMP-binding domains) are the third most abundant domain in known natural product biosynthetic pathways (Table S1). Despite the essential and varied roles of ANL enzymes, there is no database that catalogs their biosynthetic diversity. Therefore, we mined the literature, MIBiG (15), and UniProtKB (16) for ANL enzymes with known substrate specificities. We then constructed a training set of >1,700 ANL protein sequences paired with their functional class, substrate(s), kinetic data, and crystal structures if solved. As reported previously by Gulick and others, ANL superfamily enzymes in our training set were divergent at the sequence level but shared a common structural fold and core motifs including (Y/F)(G/W)X(A/T)E and (S/T)GD critical for ATP-binding and catalysis (3).

We defined nine major functional enzyme classes on the basis of having enough experimentally-characterized enzymes per class to enable classification by machine learning: short chain acyl CoA synthetases (C_2_ – C_5_, SACS), medium chain acyl-CoA synthetases (C_6_ – C_12_, MACS), long chain acyl-CoA synthetases (C_13_ – C_17_, LACS), very-long chain acyl-CoA synthetases (C_18+_, VLACS), fatty acyl-AMP ligases (FAAL), luciferases (LUC), β-lactone synthetases (BLS), aryl-CoA ligases (ARYL), and NRPS A domains (NRPS). In addition, we trained a separate model to predict enzyme substrate specificity. While prediction at the level of an individual substrate is desirable, the broad substrate specificity of some classes of ANL enzymes limited prediction to groups of chemically-similar compounds. For example, one class of LACS has demonstrated activity with fatty acids with chain lengths ranging from C_8_ – C_20_ (17). There were over 200 chemically-distinct substrates in our training set, many with just one experimental example. Therefore, we clustered substrates based on chemical similarity (Tanimoto coefficient) to identify 15 groups for broad level substrate classification (Table S2).

Previously, 34 amino acids within 8 Å of the gramicidin synthetase active site were shown to be critical for accurate prediction of NRPS A domain substrate specificity (11). Due to the high level of structural conservation between NRPS A domains and other proteins in the superfamily, we hypothesized these 34 active site residues would also be important features for ANL substrate prediction. Using an AMP-binding profile Hidden Markov Model (pHMM), we extracted 34 active site residues from our >1,700 training set sequences. Further inspection of the superfamily-wide pHMM alignment revealed the presence of a fatty acyl-AMP ligase-specific insertion (FSI) of 20 amino acids not present in other superfamily members (Fig. S1). The FSI has been suggested to inhibit the 140° domain rotation of the C-terminal domain and is critical for the rejection of CoA-SH as an acceptor molecule (6). Since the FSI was shown experimentally to be an important feature to distinguish FAAL from LACS enzymes, we extracted 20 FSI residues from each sequence and appended them to our feature vector for a total of 54 residues. Each amino acid was further encoded as 15 normalized real numbers corresponding to different physicochemical properties including hydrophobicity, volume, secondary structure, and electronic properties (11). The physicochemical properties were then used to train machine learning models to predict enzyme function and substrate (Fig. 1*A*).

**Figure 1.**
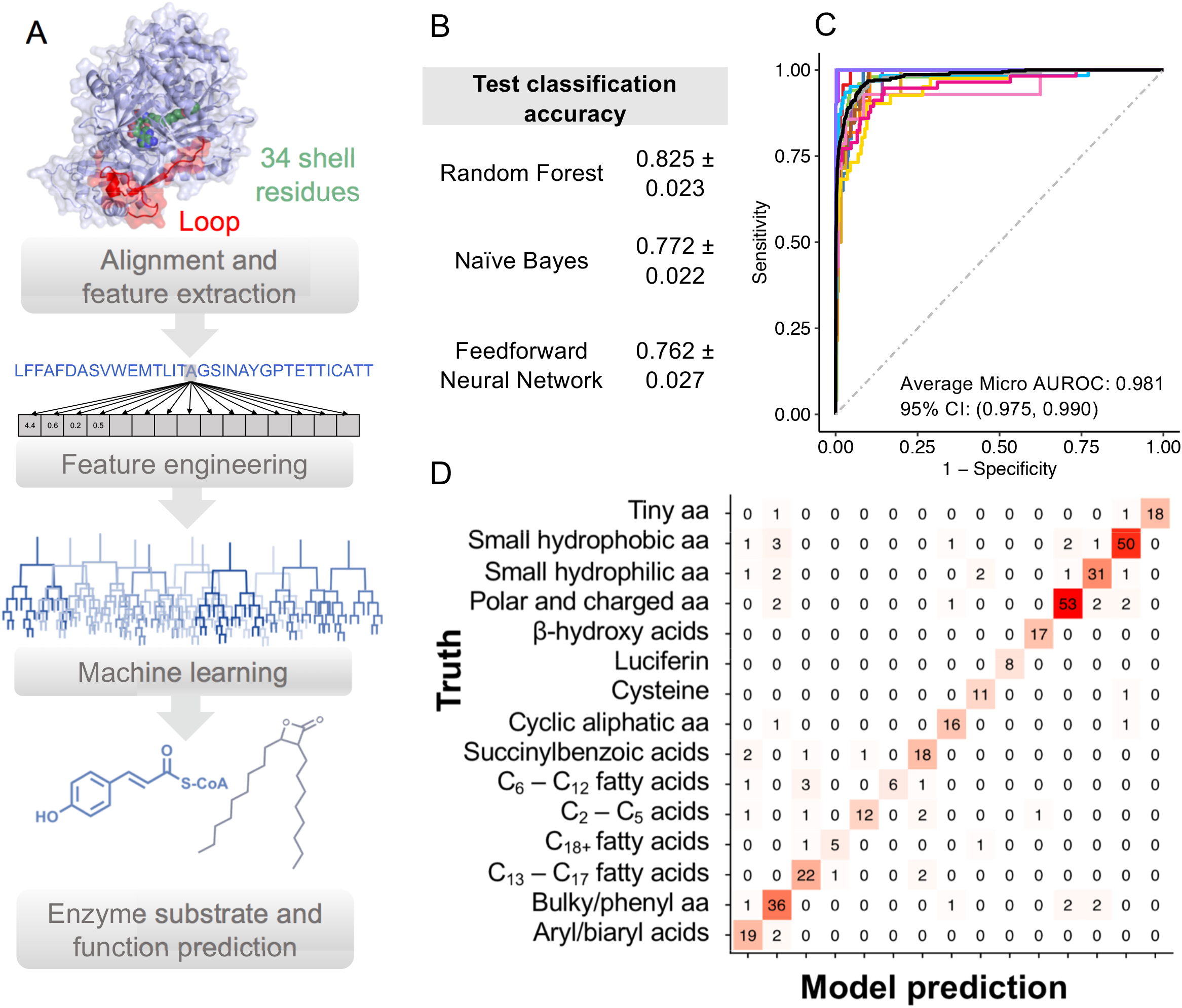
A) Workflow to predict substrate and function of adenylate-forming enzymes from protein sequence inputs. 34 active site residues (green) and FAAL-specific loop (red) residues are extracted and encoded as a vector of physiochemical properties specific to each amino acid. Separate classifiers are trained to predict substrate specificity and enzyme function. B) Hold-out test set accuracy for three different classification methods evaluated in this study C) Area under the receiver operating characteristic curve (AUROC) for substrate specificity predictions. Colors correspond to different substrates and macro (gray) and micro (black) AUROC averages. Red = aryl/biaryl acids, Green = bulky/phenyl aa, Blue = C_13_ - C_17_ fatty acids, Purple = C_18+_ fatty acids, Orange = C_2_ - C_5_ acids, Tan = C_6_ - C_12_ fatty acids, Brown = succinylbenzoic acids, Hot pink = cyclic aliphatic aa, Dark blue = cysteine, Fuchsia = luciferin, Light pink =ß-hydroxy acids, Turquoise = polar and charged aa, Goldenrod = small hydrophilic aa, Deep pink = small hydrophobic aa, Lavender = tiny aa. aa = amino acids. D) Confusion matrix of predicted vs. truth for ANL substrate specificity on hold-out test set. Predictions for functional class are presented in Figure S2.

Three different machine learning algorithms were evaluated for our classification problem: feedforward neural networks, naïve Bayes, and random forest. Random forest performed slightly better than its counterparts for both function and substrate classification problems (Fig. 1*B* and Fig. S2). Naïve Bayes also performed well, but was significantly slower than random forest in run-time. The feedforward neural network performed the worst, likely due to a relatively small number of training samples, and was also the slowest model to train. On the basis of speed and accuracy we chose to proceed with the random forest algorithm. Our best performing model achieved 82.5 ± 2.3% test set classification accuracy for substrate specificity and 83.3 ± 3.0% for functional class prediction (Fig. 1*B* and Fig.S2). The average area under the receiver operating characteristic curve (micro-average AUROC) was 0.981 for substrate group prediction and 0.978 for functional class (Fig. 1*C* and Fig. S2). Within-class accuracy was highest for the FAAL and NRPS A domains, most likely due to a larger amount of experimental data for these protein families (Fig. S2). Both of our classifiers also performed well with more specialized enzymes that accept single substrates such as the BLS and LUC classes (Fig. S2). We speculate that enzymes with specialized functions might have distinct ‘active site patterns’ that could be learned by our algorithms due to the preference of these enzymes for single substrates (e.g. D-luciferin). Our machine learning model consistently performed the worst on substrate classification for enzymes with broad substrate specificity such as the aryl/biaryl acids and C_6_ – C_11_ chain length fatty acids and aryl acids (Fig. 1*D*).

We developed a web application, AdenylPred (z.umn.edu/AdenylPred), for **<u>Adenyl</u>** ation **<u>Pred</u>**iction to make our machine learning models publicly available. Users can upload their sequences of adenylate-forming enzymes in multi-FASTA or GenBank format either as nucleotide or protein sequences. Predictions and probability scores are reported for both functional classification and substrate specificity. The entire ANL enzyme training set is also available in a searchable database format. Overall, the web app provides an interactive interface for users with little computational experience. A command-line version of the tool is also available for download for the analysis of large datasets.

### Specialized ANL enzymes may have evolved from an ancestral scaffold utilizing CoA-SH

We next used a phylogenetic approach to investigate the functional divergence of ANL superfamily enzymes (Fig. 2). Maximumlikelihood phylogenetic analysis revealed that some functional classes, including the BLS, LUC, and NRPS enzymes, formed tight monophyletic clades whereas others, mainly enzymes in the ARYL class, were dispersed throughout the tree. The wide phylogenetic distribution of the ARYL sequences indicated many ARYL enzymes were more closely related to different protein subfamilies than to each other. This observation could likely be explained by two evolutionary scenarios: 1) aryl-CoA ligase activity arose independently several times throughout ANL superfamily evolution or 2) radial divergence of the superfamily occurred from an ancestral scaffold similar to contemporary ARYL enzymes (18). To investigate these scenarios further, we used a maximum-likelihood approach for ancestral sequence reconstruction to estimate the 10 most likely sequences for the predicted ancestral protein at the root of the ANL phylogeny (19). We used AdenylPred to extract sequence features and predict function and substrate for our reconstructed ancestral proteins. AdenylPred classified 10/10 of the most likely ancestral ANL proteins as aryl-CoA ligases most likely to activate aryl and biaryl derivatives as substrates (probability score = 0.6). We also tested a maximum-likelihood ancestral reconstruction using only 34 active site residues as the seed sequences rather than full sequences and obtained similar ARYL predictions (probability score = 0.7). These findings suggest that the active sites of our reconstructed ancestral ANL proteins were most similar to contemporary ARYL enzymes.

**Figure 2.**
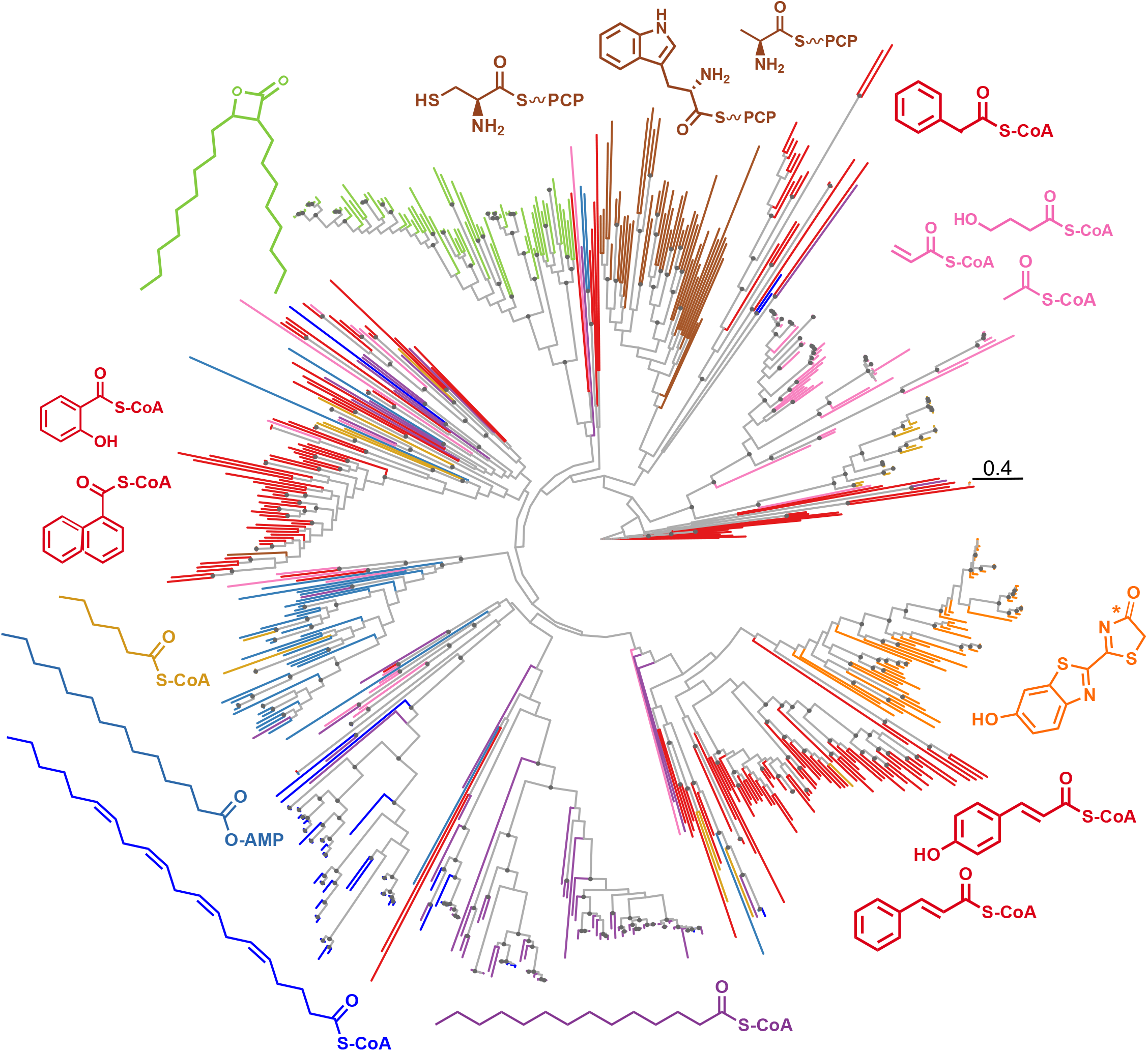
Maximum-likelihood phylogenetic tree of characterized protein sequences in the ANL superfamily computed using the Jones-Taylor-Thornton matrix-based model of amino acid substitution and colored by functional enzyme class. Some enzyme classes such as BLS, NRPS, and LUC form monophyletic clades while other sequences i.e. the ARYL class are dispersed throughout the tree, suggesting evolutionary divergence. Red = ARYL, Green = BLS, Dark blue = VLACS, Orange = LUC, Light blue = FAAL, Brown = NRPS, Purple = LACS, Pink = SACS, Gold = MACS. Gray node circles represent bootstrap support >75% at branch points. Bar, 0.4 aa substitutions per site.

### Sequence similarity networking reveals a ‘hub’ network topology

To construct a map the distribution of adenylate-forming enzymes in biosynthetic gene clusters, we applied AdenylPred to a taxonomically-diverse and representative collection of bacterial, fungal, and plant genomes. We extracted 50,064 standalone adenylate-forming enzyme sequences from candidate biosynthetic gene clusters detected using antiSMASH (20), fungiSMASH (21), and plantiSMASH (22). To visualize results, we constructed a sequence similarity network with adenylate-forming enzymes colored by their functional classification using AdenylPred (Fig. 3*A*). The sequence similarity network displayed a ‘hub-and-spoke’ topology in which most functional protein subfamilies showed higher sequence similarity with hub sequences than with any other subfamily (Fig. 3*A*). The hub nodes were mainly comprised of aryl-CoA ligases. To test the robustness of this topology with a different sequence set, we constructed a sequence similarity network from a smaller set of all AMP-binding domains from the manually-curated MIBiG database (Fig. S3). Again, we recovered the same topology with the aryl-CoA ligases at the center. Of the 48,250 full length AMP-binding hits that we analyzed with AdenylPred, the majority were predicted to be in the ARYL and NRPS classes (Fig. 3*B*). There were no predicted luciferases in the biosynthetic gene clusters which was expected since insect genomes including fireflies were not included in the dataset. Interestingly, the predicted FAALs outnumbered LACS more than tenfold (3,673 to 322). Since both FAALs and LACSs both accept long-chain fatty acids as substrates, these findings support previous reports that the majority of lipid tails in lipopeptides and other natural products may be incorporated through FAAL-mediated activity rather than CoA-activation (23). β-Lactone formation and CoA-activation of long chain fatty acids and were the least common functions likely catalyzed by ANL enzymes in biosynthetic gene clusters (Fig. 3*B*).

**Figure 3.**
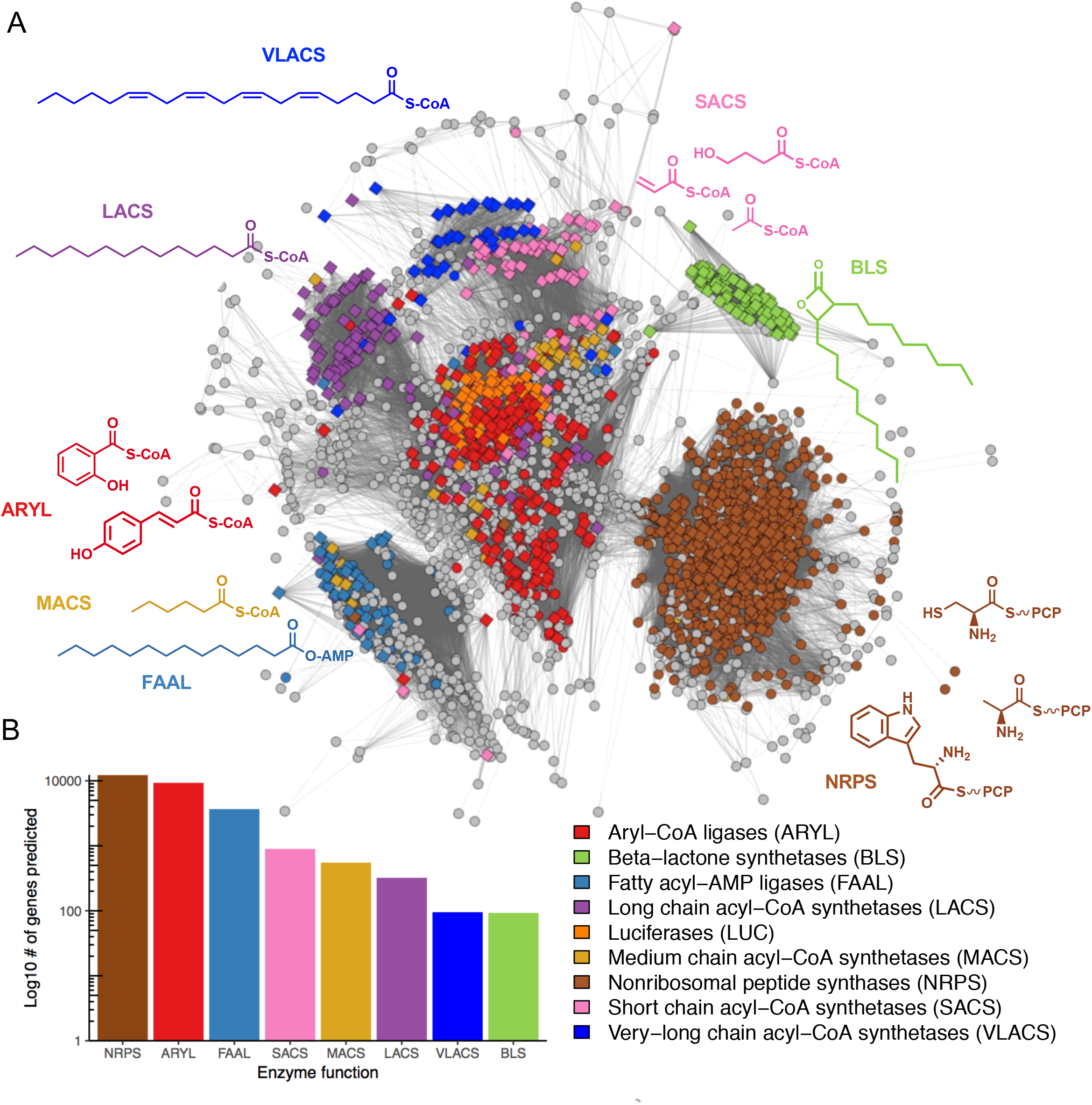
A) Sequence similarity network of all AMP-binding pHMM hits from 50,064 predicted biosynthetic gene clusters in bacterial, fungal, and plant genomes. Diamonds correspond to training set sequences and circles represent AMP-binding hits extracted from biosynthetic gene clusters. The network was trimmed to a BLAST e-value threshold of 1 x 10^-36^. Circles with a probability > 0.6. are colored by their prediction, while sequences colored gray have ‘no confident prediction.’ B) Bar plot of relative distribution of functional classes of ANL enzymes within biosynthetic gene clusters (AdenylPred prediction probability > 0.6).

### AdenylPred-guided discovery of β-lactone synthetases in biosynthetic gene clusters

β-Lactone synthetases were the most recently discovered members of the ANL superfamily and comprehensive analysis of their prevalence in natural product biosynthetic gene clusters has never been conducted. We examined AdenylPred hits for β-lactone synthetases from our collection of 50,064 candidate biosynthetic gene clusters. As expected, we detected candidate β-lactone synthetases in gene clusters responsible for the biosynthesis of known β-lactone natural products including ebelactone and lipstatin (24, 25). We also unexpectedly detected uncharacterized candidate β-lactone synthetases in >90 genomes belonging to the bacterial genus *Nocardia*. Only one β-lactone natural product, nocardiolactone, from *Nocardia* spp., has been isolated to date (26). No follow-up studies on nocardiolactone were published and the biosynthetic gene cluster was never reported resulting in nocardiolactone being termed an ‘orphan’ natural product.

We identified three flanking genes around the predicted β-lactone synthetase genes in *Nocardia* spp. that shared synteny with biosynthetic genes for the β-lactone natural product, lipstatin (Fig. 4*A*). The lipstatin cluster encodes NRPS and formyltransferase enzymes that attach an *N*-formyl leucine to the di-alkyl backbone (23). Notably, NRPS and formyltransferase genes were absent from all candidate clusters in *Nocardia* (Fig. 4*B*). We hypothesized that this minimal set of biosynthetic genes, termed *nltABCD*, encode the necessary enzymes for *Nocardia* spp. to produce a di-alkyl β-lactone product similar to lipstatin but lacking an amino acid side chain. Predicted functions of genes in the *nltABCD* operon correspond to reactions required to produce the orphan structure of the β-lactone product, nocardiolactone (Fig. 4*A*), prompting us to characterize the biosynthetic enzymes and pathway experimentally.

**Figure 4.**
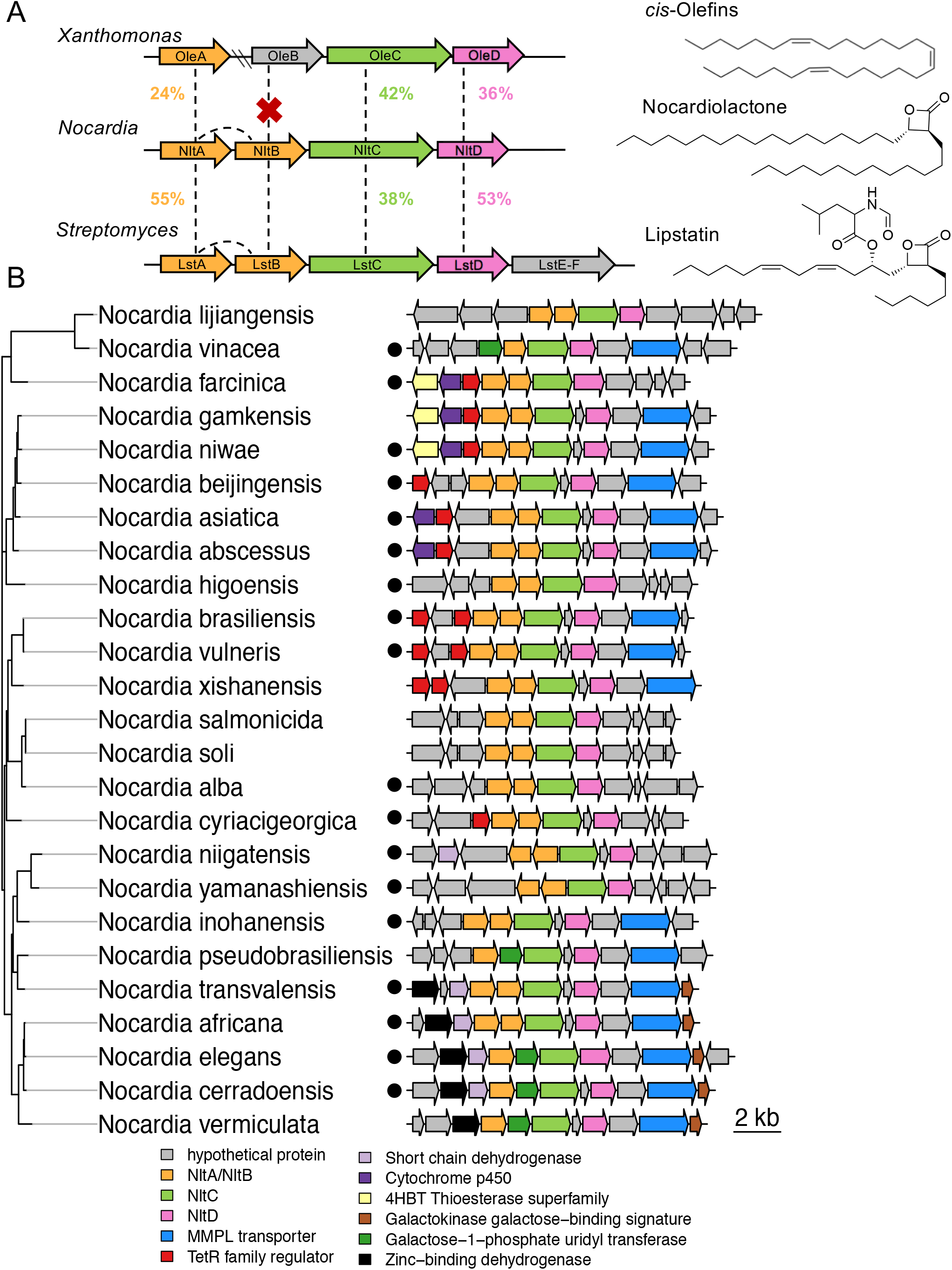
A) Synteny between published bacterial c/s-olefin and lipstatin gene clusters with the proposed nocardiolactone biosynthetic gene cluster. Percentages correspond to amino acid identity. B) Representatives of the proposed nocardiolactone biosynthetic cluster in *Nocardia.* Maximum-likelihood phylogenetic tree is based on NltC amino acid sequence distance estimated using the Jones Taylor-Thornton model of amino acid substitution. Sequences corresponding to *Nocardia* isolated from humans are designated by black circles.

### *In vitro* reconstitution of the nocardiolactone pathway links biosynthetic gene cluster to its orphan natural product

Nocardiolactone was originally isolated in 1999 from a pathogenic strain of *Nocardia brasiliensis* and other unidentified strains of *Nocardia* spp., none of which are available in public culture collections (26). Genetic manipulation in *Nocardia* is challenging therefore we opted instead to reconstitute the complete biosynthetic pathway *in vitro* by heterologously expressing and purifying individual *nltABCD* pathway enzymes. This approach gave us full control to determine the function of each pathway enzyme through biochemical analysis of intermediates and comparison to synthetic standards.

We cloned and expressed a candidate β-lactone synthetase, termed NltC, from a publicly available *N. brasiliensis* genome. NltC purified as a 60 kDa monomer. To test NltC for β-lactone synthetase activity, we incubated NltC, ATP, and MgCl_2_ with *syn*- and *anti*-2-octyl-3-hydroxydodecanoic acid diastereomers synthesized as substrate analogs (Fig. 5*A*). We observed NltC-catalyzed formation of *cis*-and *trans*-β-lactones after overnight reaction compared to no enzyme controls (Fig. S4*A*). The coupling constants were consistent with synthetic standards for *cis*- and *trans*-β-lactones of comparable chain length. AMP release is also commonly used as a readout for ANL superfamily activity. In a time-course analysis of AMP release by NltC, we observed activity with C_20_ chain length β-hydroxy acids but no activity with C_14_ length analogs above the level of the no-substrate control (Fig. 5*A*). NltC also showed weak activity with 2-hexyldecanoic acid as a substrate mimic, but not with 10-nonadecanol, suggesting adenylation of the carboxylic acid occurs rather than hydroxyl group (Fig. S4*B*). These results are consistent with the reported adenylation activity of for the β-lactone synthetase from *Xanthomonas campestris* and with the ANL superfamily (27). Overall, these results support AdenylPred-guided predictions that NltC in *N. brasiliensis* is a functional β-lactone synthetase.

**Figure 5.**
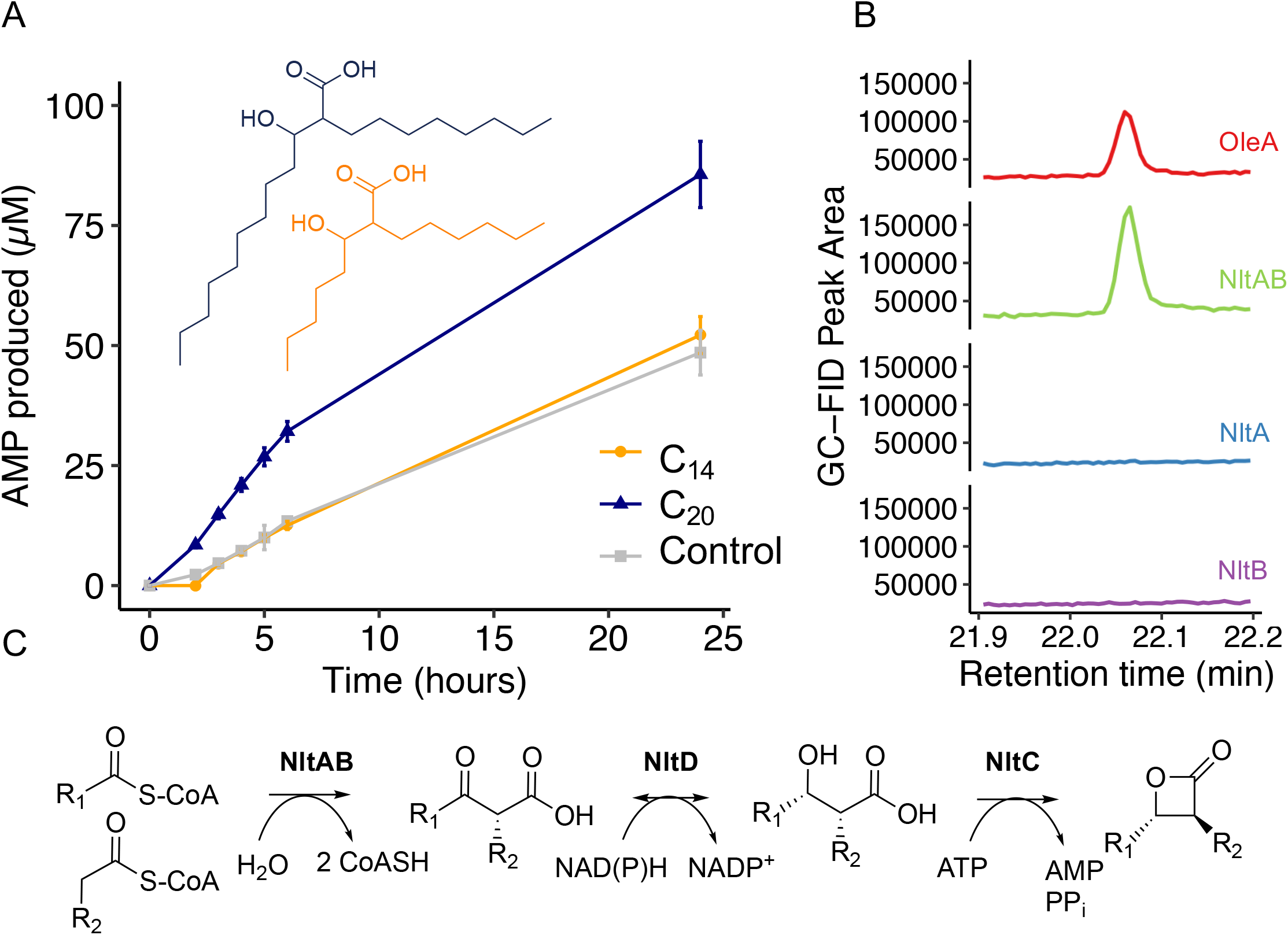
A) Time-course analysis of NltC activity with di-alkyl β-hydroxy acids with carbon backbones of length C_20_ (blue) and C_14_ (orange) compared to a no substrate control (gray). NltC prefers longer chain β-hydroxy acids (C_20_) and shows no discernable activity with C_14_β-hydroxy acids above the level of the no substrate control. B) Co-expressed NltAand NltB enzymes condense 2 myristoyl-CoAs to form 2-myristoyl-3-ketomyristic acid. The resulting ketone (14-heptacosanone) from the breakdown of 2-myristoyl-3-ketomyristic acid was observable by GC-MS. The enzymatic product of NltAB was identical to a 14-heptacosanone control produced by wild-type *X. campestrís* OleA but was not observed to be catalyzed by NltA or NltB enzymes purified individually. C) Proposed biosynthetic pathway for nocardiolactone. R_1_ = C_18_H_37_, R_2_ = C_13_H_27_.

To further characterize enzymes involved in nocardiolactone biosynthesis, we heterologously expressed and purified two upstream thiolases, NltA and NltB. Recently, homologous enzymes in the lipstatin biosynthetic pathway, LstA and LstB were shown to form a functional heterodimer to catalyze ‘head-to-head’ Claisen condensation of two acyl-CoAs (28). We hypothesized NltA and NltB might catalyze a similar reaction to form the nocardiolactone backbone (Fig. 5*B*). When heterologously expressed, NltA was mostly insoluble and formed inclusion bodies even when co-expressed with chaperones. We also attempted to purify a NltA homolog from a closely-related organism, *Nocardia yamanashiensis,* and again observed inclusion body formation. In contrast, NltB could be purified with a moderate yield (~28 mg/L of culture). When tested individually, NltA and NltB protein preparations did not display activity with any chain length (C_8_ – C_16_) acyl-CoA substrate tested. However, when NltA and NltB were co-expressed on a single plasmid we obtained soluble protein that actively catalyzed the Claisen condensation of long chain acyl-CoAs to β-ketoacids (Fig. 5*B*). Size exclusion chromatography indicated that co-expressed NltAB eluted at approximately 72 kDa, suggesting a heterodimeric state.

From homology modeling and sequence analysis we observed that the NltB sequence was truncated and lacked a Cys-His-Asn catalytic triad similar to reports for LstB (28). Although no crystal structure of LstAB is available, site-directed mutagenesis revealed a conserved glutamate in LstB (E60) was required for condensation activity (28). In the crystal structure of the physiological homodimer, OleA, from *Xanthomonas campestris*, a similarly-positioned glutamate from the β-chain was shown to enter the active site of the α-chain in the OleA homodimer and deprotonate the α-carbon to activate it (29). Based on homology modeling and structural alignments with OleA, NltB_E57_ may be similarly poised to act as a nucleophile in the NltA active site (Fig. S5*A*). We used site-directed mutagenesis to mutate the NltB glutamate (E57) to either alanine or glutamine. Claisen condensation activity was abolished in NltAB_E57A_ and NltAB_E57Q_ mutants compared to wild-type NltAB (Fig. S5*B*). Taken together, these results suggest NltA and NltB may form a functional heterodimer with NltB_E57_ required to catalyze the Claisen condensation of two fatty acyl-CoA substrates. Crystallographic studies are required to investigate LstAB and NltAB further.

The final enzyme in the nocardiolactone cluster, NltD, belongs to the short chain reductase superfamily. NltD purified as a 78 kDa fusion protein with a maltose binding protein tag. NltD has an N-terminal conserved nucleotide binding motif (Rossman fold) and a SX_*n*_YXXXK catalytic triad characteristic of short chain reductase superfamily members (30). NltD shares 53% amino acid identity with the lipstatin reductase (LstD) and 36% identity with *X. campestris* OleD, a 2-alkyl-3-ketoalkanoic acid reductase involved in olefin biosynthesis (Fig. 4*A*). Since studies on OleD demonstrated 2-alkyl-3-ketoalkanoic acids are unstable, we monitored the reaction in reverse with 2-alkyl-3-hydroxyalkanoic acid substrates using a spectrophotometric assay for NADPH formation (30). Purified NltD catalyzed NADP^+^-dependent conversion of 2-alkyl-3-hydroxyalkanoic acid to 2-alkyl-3-ketoalkanoic acid with both C_20_ and C_14_ di-alkyl β-hydroxy acid substrates at a rate similar to *X. campestris* OleD (Fig. S6).

We next reconstituted the entire pathway to produce nocardiolactone-like analogs by combining purified pathway enzymes with decanoyl-CoA precursors, NADPH, ATP, and MgCl_2_ (Fig. 5*C* and Fig. S7). Due to instability and loss of activity of purified NltAB over time, we substituted the functionally-equivalent and stable homodimer, OleA, from *X. campestris* (31). After overnight incubation, we observed formation of a di-alkyl β-lactone natural product (Fig. S7). Based on *in vitro* evidence, we propose nocardiolactone biosynthesis is initiated via ‘head-to-head’ Claisen condensation of fatty acyl-CoA substrates catalyzed by a heterodimeric interaction between NltA and NltB. NltD then reduces the dialkyl β-keto acid to a di-alkyl β-hydroxy acid in an NADPH-dependent manner. Finally, intramolecular ring closure of a β-hydroxy acid to a β-lactone is catalyzed by the ATP-dependent β-lactone synthetase NltC (Fig. 5*C*). Overall, these results link the *nltABCD* biosynthetic gene cluster in *Nocardia brasiliensis* to the orphan natural product, nocardiolactone.

### The nocardiolactone gene cluster is enriched in human pathogens

With the biosynthetic gene cluster established, we next probed the taxonomic distribution and abundance of the nocardiolactone pathway in *Nocardia* genomes. We used AdenylPred to identify putative β-lactone synthetases and detected *nltC* homologs with flanking *nltABD* genes in 94 out of 159 complete *Nocardia* genomes in the PATRIC database (32). Notably, the strict *nltABCD* cluster was not detected in any closely related genera such as *Rhodococcus, Streptomyces*, or *Mycobacterium*, suggesting nocardiolactone biosynthesis may be specific to the genus *Nocardia*. We observed the nocardiolactone gene cluster was more prevalent among strains of pathogenic clinical isolates from human patients than in strains isolated from other sources (Fig. 4). The complete *nltABCD* cluster was detected in 68% of genomes of distinct species of human pathogenic *Nocardia* relative to 27% isolated from non-human sources (p-value: 0.002, Fisher’s exact test). We also queried a separate private database comprised of 169 clinical isolates of *Nocardia* from human patients (Pidot, Stinear, *in prep*) and recovered a similar proportion of complete nocardiolactone cluster hits among clinical isolates (112/169, ~66%). Although only correlative, the enrichment warrants further research and suggests a potential link between the nocardiolactone cluster and *Nocardia* pathogenicity.

## DISCUSSION

Based on our ancestral reconstruction, we propose ancient ANL enzymes had an active site most similar to contemporary enzymes that activate aryl and biaryl acids with CoA. This hypothesis is supported by the unusual hub topology of the ANL network. Many other enzyme superfamilies do not have this topology and instead tend to show patterns of sequential functional divergence (33, 34). However, Babbitt and colleagues detected a similar hub topology in the nitroreductase superfamily and provided multiple lines of evidence supporting that the topology indicated radial divergence of the superfamily from a minimal flavin-binding scaffold (18). Our results also suggest ANL sequences may have undergone divergent evolution towards more specialized functions from ancestral enzymes with CoA-ligase-like scaffolds.

The proposed evolutionary trajectory of the ANL superfamily is supported by experimental evidence for CoA-ligase activity in many extant ANL enzymes that primarily perform other functions. For example, Linne et al. tested the CoA-ligase activity of five different NRPS A domains (35). Surprisingly, all the NRPS A domains were also able to synthesize acyl-CoAs *in vitro*. Enzymatic CoA-ligase activity of the NRPS A domains varied proportionally to their evolutionary similarity with a native acyl-CoA synthetase, suggesting that greater sequence divergence resulted in more specialized NRPS A domain activity and reduced bifunctionality (35). Firefly luciferases were also demonstrated to be bifunctional as CoA-ligases (36) and luciferase activity was conferred to an acyl-CoA ligase from a non-luminescent click beetle by just a single point mutation (37). Arora et al. showed FAAL enzymes likely lost their CoA-ligase activity due to the FSI. Indeed, deletion of the FSI conferred acyl-CoA ligase activity in FAAL28 from *Mycobacterium tuberculosis* (6). The fact that simple mutations can revert many enzymes back to a CoA-ligase state supports the hypothesis that the ANL superfamily arose from an ancestral enzyme using CoA-SH as an acceptor molecule. However, we cannot rule out the alternative hypothesis of CoA-ligase activity arising independently several times in the ANL superfamily. Overall, ancestral reconstruction coupled with AdenylPred analysis yielded new insights into the evolutionary structure-function relationships among adenylate-forming enzymes.

Within the ANL superfamily, the β-lactone synthetases were most recently discovered and the extent of their role in natural product biosynthesis remained poorly understood (8, 27). We used AdenylPred to detect >90 β-lactone synthetases in uncharacterized biosynthetic gene clusters which is significantly more than the eight known biosynthetic gene clusters for β-lactone natural products reported to date (38). The disparity between the number of predicted β-lactone biosynthetic gene clusters and known β-lactone natural products has several possible explanations. One reason may be limited discovery due to the reactivity and thermal instability of β-lactones. β-lactones are strained rings that can rapidly hydrolyze in aqueous solutions (38) or thermally decarboxylate, thus hampering their detection by common analytical methods such as GC-MS (8). It is also plausible many biosynthetic gene clusters with β-lactone synthetases are not expressed under normal laboratory conditions. Another explanation is the undetected role of β-lactones as intermediates in the biosynthesis of other chemical moieties (8). Chemists have long referred to β-lactones as ‘privileged structures’ for the total synthesis of compounds with a variety of functional groups including β-lactams, gamma-lactones, and alkenes (38, 39). Our findings suggest microbes might also use β-lactones as intermediates since we detected a number of β-lactone synthetase hits in gene clusters known to make natural products without final β-lactone moieties such as polyunsaturated fatty acids. Indeed, β-lactone synthetases in *oleABCD* gene clusters were recently linked to production of the final alkene moiety in the biosynthesis of a C31 polyunsaturated hydrocarbon product (40). Such cases, if more widespread, may have also escaped detection because most of the predicted gene clusters with β-lactone synthetases were not detected by tools like antiSMASH (20) until recently, prohibiting the discovery of such biosynthetic pathways through genome mining approaches. The recent feature implementation in antiSMASH to detect likely β-lactone moieties now enables further research into the role of β-lactones as intermediates in the biosynthesis of hydrocarbons and other secondary metabolites.

Among the predicted gene clusters with β-lactone synthetase homologs, we detected a conserved cluster in *Nocardia* which we linked to the orphan β-lactone natural product, nocardiolactone. Numerous biosynthetic gene clusters have been detected in *Nocardia* spp., and many genomes from *Nocardia* have as many type I polyketide synthases and NRPS gene clusters as *Streptomyces* genomes (41). Despite this, only a small number of biosynthetic gene clusters in *Nocardia* have been experimentally-verified. Recently, the biosynthetic gene cluster for the nargenicin family of macrolide antibiotics was discovered in human pathogenic strains of *Nocardia arthriditis* (42). Khosla and colleagues also reported on a unique class of orphan polyketide synthases in the genomes of *Nocardia* isolates from human patients with nocardiosis (43). Based on the conservation of this gene cluster in clinical isolates from nocardiosis patients, Khosla proposed the product might play a role in *Nocardia* pathogenicity in human hosts. Similarly, we found the nocardiolactone gene cluster was significantly more abundant in the genomes of human pathogenic strains compared to isolates from non-human sources. The enrichment supports the hypothesis that nocardiolactone could play a role in the pathogenicity of *Nocardia* however further *in vivo* studies are required.

Nocardiolactone was first isolated from the mycelia of *Nocardia* spp. rather than the fermentation broth, suggesting the compound is cell-associated (26). The long, waxy di-alkyl tails of nocardiolactone resemble mycolic acids and would likely embed in the cell membrane. The cell wall composition varies between different species of *Nocardia* but is known to consist primarily of trehalose dimycolate and several other unidentified hydrophobic compounds (44). Studies on *Nocardia* virulence found that cell-surface composition was a critical determinant for attachment and penetration of host cells (45). Cell wall-associated lipids in *N. brasiliensis* were also shown to induce a strong inflammatory response (44). The role of hydrophobic natural products in *Nocardia* pathogenicity and infection remains a rich and untapped direction for future research.

In summary, we conducted the first global analysis of adenylate-forming enzymes in >50,000 candidate biosynthetic gene clusters from all domains of life. Our machine learning approach yielded evolutionary insights into ANL superfamily divergence from a core scaffold similar to contemporary aryl-CoA ligases towards enzymes with more specialized functions such as β-lactone formation. AdenylPred analysis also detected >90 β-lactone synthetases in gene clusters in *Nocardia* spp. that were enriched in the genomes of human pathogens. Through *in vitro* pathway reconstitution, we were able to link this gene cluster family to the orphan natural product nocardiolactone. These findings demonstrate how machine learning methods can be used to pair gene clusters with orphan secondary metabolites and advance understanding of natural product biosynthesis.

## EXPERIMENTAL PROCEDURES

### Sequence similarity network

Candidate biosynthetic gene clusters from >24,000 bacterial genomes in the antiSMASH database version 2 (46) were combined with pre-calculated plantiSMASH output (22) and results from fungiSMASH analysis of 1,100 fungal genomes (accession numbers available at https://github.com/serina-robinson/adenylpred_analysis/). The AMP-binding pHMM (PF00501) was used to query all genes in the dataset with default parameters, returning 213,993 significant hits. Since the distribution and identity of NRPS adenylation domains have already been analyzed in detail by Chevrette et al. (12), we opted to analyze only standalone AMP-binding pHMM hits. All sequences with a condensation domain (PF00668) in the same coding sequence were filtered out, leaving 71,331 ‘standalone’ AMP-binding HMM hits. Of these, partial sequences (less than 150 amino acids in length) were removed for a total of 63,395 sequences. Due to computational limitations of visualizing large networks, the sequences were clustered using CD-HIT (47) with a word size of 2 and 40% sequence similarity cutoff to yield 2,344 cluster representatives. Cluster representatives were combined with the AdenylPred training set sequences to observe their relation to sequences with known specificity. A sequence similarity network was constructed by calculating pairwise BLAST similarities between all sequences. The network was examined over a comprehensive range of e-value cutoffs and visualized using the ‘igraph’ package (48).

### Training set construction

The AdenylPred training set was pulled and manually-curated from three databases: UniProtKB, MIBiG, and the most up-to-date NRPS A domain training set from SANDPUMA (12). AMP-binding enzymes (PF00501) in the UniProtKB database that had experimental evidence at the protein level were extracted and linked to their substrate through literature mining and manual verification. The MIBiG database was queried with the AMP-binding HMM (PF00501). MIBiG sequences were extracted and linked to their product and substrate when reported in the literature. The SANDPUMA NRPS A domain dataset (12) was randomly down-sampled with stratification by substrate class to balance the training set classes for substrate and enzyme function prediction. Training set sequences were grouped into 15 substrate groups and 9 enzyme functional classes.

### Machine learning methods

Protein sequences in the training set were aligned with the AMP-binding HMM using HMMAlign (49). 34 substrate residues within 8 Å of the active site were extracted and encoded as tuples of physicochemical properties as described by Röttig et al. (11). The FSI was extracted, one-hot encoded, and concatenated to the vector of active site vector properties for a total of 585 sequence features. Three different machine learning algorithms were trained for multiclass classification: random forest, naïve Bayes, and feedforward neural networks. The data were split with stratified sampling into 75% training and 25% test sets. Tuning parameters for all models were adjusted by grid search using 10-fold cross validation repeated with 5 iterations. The confidence of random forest predictions can be assessed using a nonparametric probability estimation for class membership calculated as a value between 0 and 1 (50). Based on the distribution of prediction probabilities, we set an empirical threshold for prediction confidence of 0.6 (60%), below which all substrates are listed as ‘no confident result’ although the best prediction is still provided to the user.

### AdenylPred availability

The web application is available at z.umn.edu/adenylpred(shortened url) or https://srobinson.shinyapps.io/AdenylPred/. The command-line version of the tool is available at https://github.com/serina-robinson/adenylpred.

### Phylogenetic analysis and ancestral reconstruction

Training set sequences were aligned were aligned using HMMAlign (49) and terminal ends of the alignment were trimmed. The phylogeny of the entire training set was estimated using RAxML (51) with the Jones-Taylor-Thornton matrix-based model of amino acid substitution and a discrete gamma model with 20 rate categories. For ancestral sequence reconstruction, training set sequences were redundancy filtered to 40% amino acid identity with a word size of 2 with CD-HIT (47). FastML was used to reconstruct the most likely ancestral sequences for internal nodes of the tree (19). AdenylPred was then used to predict the functional class and substrate of the root ancestral sequence in the ANL superfamily.

### HPLC analysis of NltC activity

HPLC analysis of ATP, ADP, and AMP was conducted using an Agilent 1100 series instrument with a C_18_ eclipse plus (Agilent) column mounted with a C_18_ guard column. Reactions were carried out in glass HPLC vials with a total volume of 500 μL in assay buffer (50 mM Tris base adjusted to pH 8.0 with HCl). Reactions were initiated with the addition of 0.5 μM enzyme (15 μg) to 100 μM of ATP, 100 μM of substrate, and 2% EtOH originating from the substrate stocks. Separation of ATP, ADP, and AMP was observed after 9 min under isocratic conditions with 95% 100 mM H2KPO4 (pH 6.0 with KOH) and 5% methanol while monitoring at 259 nm.

### Cloning, Expression, and Purification of Nocardiolactone Biosynthetic Enzymes

NltA (WP_042260942.1), NltB (WP_042260944.1), NltC (WP_042260945.1) and NltD (WP_042260949.1) from *Nocardia brasiliensis* and NltA from *Nocardia yamanshiensis* (WP_067710538.1), were codon-optimized for *E. coli* and synthesized by Integrated DNA Technologies (IDT, USA). NltC was cloned into a pET30b+ vector with NdeI and HindIII restriction sites with a C-terminal 6x-His tag. NltD was cloned into a modified pMAL-c5x vector with a tobacco etch virus (TEV) protease cut site added at NdeI and HindIII restriction sites and a N-terminal 6x-His tag and expressed as a fusion with maltose binding protein. NltA from *N. yamanashiensis* was cloned into a pET28b+ at NdeI and XhoI restriction sites. NltA and NltB from *N. brasiliensis* were cloned both individually as in the case of *N. yamanashiensis* and combined with a ribosome binding site-like sequence (tttgtttaactttaagaaggaga) inserted into a single pET28b+ vector with a N-terminal 6x-His tag. Accession numbers and codon-optimized plasmid sequences are available in the Supporting Information. Constructs were cloned by Gibson assembly into DH5α cells and verified by Sanger sequencing. Sequence-verified plasmids were transformed into BL21 (DE3) cells (NEB). Starter cultures (5 mL) were grown in Terrific Broth overnight at 37° C with kanamycin selection. 1-Liter cultures with 75 μg/ml kanamycin were grown to an optical density of 0.5 at 37° C, induced by addition of 1 mL isopropyl β-D-1-thiogalactopyranoside (IPTG, 1 M stock) and further incubated for 19 h at 15 °C. Induced cells were harvested at 4000 x g with a Beckman centrifuge and frozen at −80 °C. Cell pellets were resuspended in 10 mL of buffer containing 500 mM NaCl, 20 mM Tris base, and 10% glycerol at pH 7.4. Elution buffer for the nickel column was the same but with the addition of 400 mM imidazole. NltD purification buffer required the addition of 0.025% Tween 20. Cell pellets were thawed on ice, lysed with 2–3 cycles in a French pressure cell (1500 pounds per square inch) and centrifuged for 60 min at 17,000 x g. Supernatants were filtered through a 0.45 μM low protein binding filter (Corning, USA), loaded into a GE Life Sciences ÄKTA fast liquid protein chromatography system and injected onto a GE Life Sciences HisTrap HP 5 ml column. After washes to remove nonspecifically bound proteins, His-tagged proteins were eluted with a stepwise gradient of 5, 10, 15, and 80% elution buffer over 2.5 column volumes at 1 ml/min and collected in 2 ml fractions. Protein concentration was determined by the method of Bradford (52) using the Bio-Rad (USA) Protein Assay Dye Reagent Concentrate and a standard curve prepared from a 2 mg/mL bovine albumin standard (Thermo Scientific). The desired protein fractions were pooled, analyzed by SDS-PAGE, flash frozen in liquid N2 and stored at −80°C.

### GC-MS

Separation and identification of metabolites was accomplished by GC-MS (Agilent 7890a & 5975c) equipped with a 30 m x 0.25 mm i.d. x 0.25 μm DB-1ms capillary column with outflow split to flame ionization and MS detectors. The formation of the unstable β-keto acid catalyzed by OleA and NltAB could be observed as its ketone breakdown product by GC-MS as published previously (31). Olefins from the complete thermal decarboxylation of β-lactone products were detected without derivatization by comparison to synthetic β-lactone standards described previously (8, 27). β-hydroxy acid required methylation of the carboxylic acid group by diazomethane for detection by GC-MS. Ethereal alcoholic solutions of diazomethane were prepared from *N*-methyl-*N*-nitroso-*p*-toluenesulfonamide (Sigma-Aldrich). All samples were extracted with tert-methyl butyl ether and mixed with 50 μL of diazomethane solution. One microliter of each sample was injected into the injection port (230 °C). The 25 min program was as follows: hold 80 °C for 2 min; ramp linearly to 320 °C for 20 min; hold 320 °C for 3 min.

### Verification of NltC β-lactone synthetase activity by ^1^H-NMR

Reaction mixtures were set up in separatory funnels with 1 mg NltC, 20 mg ATP, and 30 mg MgCl_2_ • 6H_2_O in 100 mL of 200 mM NaCl and 20 mM NaPO_4_ buffer (pH 7.4). Reactions were initiated with the addition of 1.5 mL of 2-octyl-3-hydroxydodecanoic acids dissolved in EtOH (1.0 mg/mL stock). 10μL of 1-bromonaphthalene (0.5 mg/mL stock) was added to each reaction mixture as an internal standard. Reactions were allowed to run for 24 hours before 3 successive extractions were performed with 10 mL, 5 mL, and 5 mL of dichloromethane. Samples were evaporated at room temperature before solvation in CDCl_3_ for ^1^H-NMR (400 MHz). Figure S4*A* shows the ^1^H-NMR spectra of 2-octyl-3-hydroxydodecanoic acids allowed to react overnight with NltC compared to a no enzyme control. Chemical shifts for synthetic 2-octyl-3-hydroxydodecanoic acid starting materials and *cis*- and *trans*-3-octyl-4-nonyloxetan-2-one products were reported by Christenson et al. (8).

### NltD activity assay

Activity of NltD/OleD by NADPH-dependent consumption of 2-alkyl-3-ketoalkanoic acid was monitored using the method described by Bonnett et al. (30). The reaction was measured in reverse since the β-keto acid substrate for NltD/OleD homologs was previously shown to be unstable and undergoes rapid decarboxylation to a ketone (30). Briefly, progress was tracked by the change in absorbance at 340 nm by the formation or consumption of NADPH (ε_340_= 6,220 M^-1^ cm^-1^) in UV-transparent 96 well plates (Greiner, Sigma-Aldrich) measured using a SpectraMax Plus microplate reader (Molecular Devices).

### General synthetic procedures

Synthesis and purification of racemic 2-octyl-3-hydroxydodecanoic acid was performed as described previously (8, 53) via α-carbon deprotonation of decanoic acid by lithium diisopropylamide (LDA) followed by the addition of decanal to form 2-octyl-3-hydroxydodecanoate. An identical synthetic method was used to synthesize a racemic diastereomeric mix of 2-hexyl-3-hydroxyoctanoic acid from hexanal and octanoic acid as described by Robinson et al. (27).

## Data Availability

Scripts and raw data for analyses and figures presented in this manuscript are available on GitHub: https://github.com/serina-robinson/adenylpred_analysis. A searchable database for the training set is also available at z.umn.edu/adenylpred.

## Author Contributions

Conceptualization, S.L.R., M.H.M., and L.P.W.; Methodology, S.L.R., B.R.T. M.H.M., and L.P.W.; Software, S.L.R. and B.R.T.; Validation, S.L.R., B.R.T., S.J.P., and M.D.S.; Formal Analysis, S.L.R. and B.R.T.; Investigation, S.L.R., S.J.P. and M.D.S.; Data Curation, S.L.R. and B.R.T.; Writing – Original Draft, S.L.R.; Writing – Review & Editing, S.L.R., B.R.T., S.J.P, T.P.S., M.H.M., and L.P.W.; Supervision – M.H.M. and L.P.W.

## Acknowledgements

We acknowledge Satria Kautsar for assistance in extracting candidate biosynthetic gene clusters. Aalt-Jan van Dijk is recognized for his insightful discussions on machine learning. Kelly Aukema is acknowledged for manuscript edits and discussion. We are grateful to Jorge Navarro Muñoz for providing the output from fungiSMASH analysis.

## Funding

S.L.R. is supported by the National Science Foundation Graduate Research Fellowship under NSF grant number 00039202 and a Graduate Research Opportunities Worldwide (GROW) fellowship to the Netherlands supported by the NSF and the Netherlands Organization for Scientific Research (NWO) grant number 040.15.054/6097.

## Conflict of Interest

M.H.M. is on the scientific advisory board of Hexagon Bio.

## Notes

#### Summary of Updates

Updated references, minor grammatical changes, and links to code repos for reproducibility of figures and analysis

https://github.com/serina-robinson/adenylpred_analysis

