## Supporting Information for "Global analysis of adenylate-forming enzymes reveals β-lactone biosynthesis pathway in pathogenic *Nocardia*"

#### Table of Contents

#### Supporting Table and Figures

**Table S1.** Protein domains in the MIBiG database

**Table S2.** Substrate specificity groups

**Figure S1.** FAAL multi-sequence alignment

**Figure S2.** Functional group machine learning classification

**Figure S3.** MIBiG protein sequence similarity network

**Figure S4.** NltC $^1$H-NMR and AMP release assay

**Figure S5.** NltA and NltB homology modeling and site-directed mutagenesis of

**Figure S6.** NltD NADPH assay

**Figure S7.** *In vitro* reconstitution of nocardiolactone biosynthetic pathway

**Supporting File S1.** NltA and NltB codon-optimized gene constructs

**Supporting File S2.** NltC and NltD codon-optimized gene constructs

**Table S1.** Abundance of protein domains in the MIBiG database (1). ANL superfamily (AMP-binding, PF00501) domains are the third most abundant domain. *Domain count is the total number of domains for a given protein family detected from an analysis of 1,796 validated biosynthetic gene clusters in MIBiG at the time of analysis (March 2019).

| Description | PFAM ID | Domain count* |
|---|---|---|
| Ketoacyl synthase-like | PF00109; PF02801; PF16197 | 10711 |
| PP-binding | PF00550 | 6843 |
| AMP-binding | PF00501 | 3123 |
| Condensation | PF00668 | 3077 |
| Ketoredutase | PF08659 | 2736 |
| Acyltransferases | PF00698 | 2719 |
| AMP-binding C-terminal domain | PF13193 | 2367 |
| Polyketide synthase - dehydratases | PF14765 | 1884 |
| Cytochrome p450 | PF00067 | 1070 |
| ABC transporter | PF00005 | 827 |
| Thioesterase | PF00975 | 801 |
| Major facilitator superfamily | PF07690 | 546 |
| Alcohol dehydrogenase, N-terminal | PF08240 | 489 |

**Table S2.** Fifteen substrate specificity groups for adenylate-forming enzymes with representative substrates in each group. aa = amino acid.

| Substrate group name | Representative substrates |
|---|---|
| Polar and charged aa | serine, glutamine, glutamate, arginine, aspartate, diaminobutyrate, ornithine, 2-oxoisovalerate |
| Small hydrophobic aa | valine, isoleucine, 2-aminobutyric acid, norcoronamic acid, 2-hydroxyisovalerate, valinol, alpha-ketoisocaproic acid, alpha-ketoisovalerate, 2-hydroxyisocaproate |
| Bulky/phenyl aa | trypotophan, tyrosine, phenylalainine, histidine, lysine, hydroxyphenylglycine, kynurenine |
| Small hydrophilic aa | threonine, diazinane-3-carboxylic acid |
| $C_{13}$ – $C_{17}$ fatty acids | laurate, tridecanoate, myristate, pentadecanoate, palmitate, heptadecanoate |
| Ary/biaryl acids | salicylate, 2,3-dihydroxybenzoate, anthranilate, benzoate, naphthoate xanthenurate, quinolate |
| Succinylbenzoic acids | coumarate, cinnamate, vanillate, caffeate, ferulate |
| $C_2$ – $C_5$ acids | acetate, acetoacetate, acrylate, 3-hydroxypropanoate, butyrate, crotonoate, malonate, methylmalonate, oxalate, valerate, 2-ketoisovalerate |
| β-hydroxy acids | 3-hydroxy-2-octyldodecanoate |
| Cyclic aliphatic aa | proline, pipecolate |
| Tiny aa | glycine, glycolate |
| $C_{18+}$ fatty acids | stearate, oleate, docosanoate, tetracosanoate, nonadecanoate, arachidonate, eicosapentanoate, linoleate |
| Luciferin | 4,5-Dihydro-2-(6-hydroxy-2-benzothiazolyl)-4-thiazolecarboxylate |
| $C_6$ - $C_{12}$ fatty acids | hexanoate, octanoate, hydroxyoctanoate, nonanoate, decanoate |
| Cysteine | cysteine |

```
                                                  330       340       350       360       370     380
FAA32_MYCMM_Mycobacterium_marinum_FAAL          FEPYGLRETAVKPSYGLAEATLFVSTTPMDEVPTVIHVDRDELNKQRFVEVAA....DAP
FAA26_MYCTU_Mycobacterium_tuberculosis_FAAL     FAPYNLSPTAIRPSYGLAEATLYVAAPEAGAAPKTVRFDYEQLTAGQARPCGT....DGS
Q5ZTD3_LEGPH_Legionella_pneumophila_FAAL        FKEFGFRKEAFYPCYGLAEATLLVTGGTPGSSYKTLTLAKEQFQDHRVHFADD....NSP
AIW82280.1_Cylindrospermum_alatosporum_FAAL     FAAYGFRREAFYPGYGMAEATLLISGGLRKAAPVVLSVAGSNLEQNRVLVTTD....EQE
CAD32910.2_Actinoplanes_friuliensis_FAAL        FAAAGLRPEAVTPSFGMAEATLFVSGAPD.TPFVIRHADTDLLQRNEFRPATG.....DR
FAA26_MYCBO_Mycobacterium_bovis_FAAL            FAPYNLSPTAIRPSYGLAEATLYVAAPEAGAAPKTVRFDYEQLTAGQARPCGT....DGS
AEA60663.1_Burkholderia_gladioli_FAAL           FARCGFRAEAFGPCYGLAEATVFVSGKVDGAAPAFVELDQDGLARGEVREIAA....DHP
A0A0H2VDD9_ECOL6_Escherichia_coli_FAAL          FRQVNFDNKTFMPCYGLAENALAVSFSDEASGVVVNEVDRDILEYQGKAVAPG...AETR
A0R618_MYCS2_Mycobacterium_smegmatis_FAAL       FGPFGFPPKAIKPSYGLAEATLFVSTTPSAEEPKIITVDRDQLNSGRIVEVDA....DSP
AFV96135.1_Cylindrospermum_licheniforme_FAAL    FRDCGFKENFFYPCYGMAEATLFISGGFKQSPPIVKCLQAQTLKENIIVDVTHPTPSSQE
AMB48444.1_Nostoc_sp._FAAL                      FRDCGFQEKFFYPCYGMAEATLFTSGGFKQSPPIVKWVQTKALEENKIVDVTHPNPSSQE
AQX77690.1_Nodularia_sp._FAAL                   FAECGFRKEAFYPCYGMAETTLFVSGGLKTSLPVLYELEAAALEE.NLVVAAE....SKE
AAS98774.1_Lyngbya_majuscula_FAAL               FEPFGFRRDALYPAYGLAEATLLVSTKKHGEKARVLTLAAEALEKNRIVVVSP..AEKEQ
B4E7B5_BURCJ_Burkholderia_cenocepacia_ARYL      MGIDAVDIYGLSEVMGPGVASECVETK..................DGPTIWED.......
F3Y661_STIAU_Stigmatella_aurantiaca_ARYL        T...GVD...IIDGIGCTEN..........................FHIFISN.....RP
PQSA_PSEAE_Pseudomonas_aeruginosa_ARYL          ....GLE...ICDGIGATEV..........................GHVFLAN.....RP
FAC13_MYCTU_Mycobacterium_tuberculosis_LACS     ....NIE...VVQGYALTESCG........................GGTLLLS.....ED
LUCI_LUCCR_Luciola_cruciata_LUCIFERASE          .....FNLPGVRQGYGLTETT.........................SAIIITP......E
LUCI_PHOPY_Photinus_pyralis_LUCIFERASE          .....FHLPGIRQGYGLTETT.........................SAILITP......E
Q5UFR2_9COLE_Lampyris_turkestanicus_LUCIFERASE  .....FKLPGIRQGYGLTETT.........................SAIIITP......E
ACS2A_HUMAN_Homo_sapiens_MACS                   T...GLD...IRESYGQTET..........................GLTCMVS.....KT
A8LRC0_DINSH_Dinoroseobacter_shibae_SACS        L......GVPVIDHWWQTETGWAIAANP..................MGIEHLP.......
ACS1_YEAST_Saccharomyces_cerevisiae_SACS        I...GKNEIPIVDTYWQTESGSHLVTP...................LAGGVTP.......
ACSA_SALTY_Salmonella_typhimurium_SACS          I...GKEKCPVVDTWWQTETGGFMITP...................LPGAIE........
BAIB_CLOSV_Clostridium_scindens_VLACSBILE       .....LGPEKIYEMYSMTECIGLTCIR...................GDEWVKH.......
DLTA_BACCR_Bacillus_cereus_NRPS                 .....FPKATIMNTYGPTEATVAV......................TGIHVTE....EVL
DLTA_BACSU_Bacillus_subtilis_NRPS               .....FPKAKIFNTYGPTEATVAV......................TSVEITN....DVI
DLTA_STRP6_Streptococcus_pyogenes_NRPS          .....FPSAKIINAYGPTEATVAL......................SAIEITR....EMV
```

**Figure S1**. Fatty acyl AMP-ligase (FAAL) specific insertion (cyan) is conserved across enzymes with FAAL-function characterized to date but absent in other ANL superfamily members. The structural alignment was generated using DECIPHER (2) and rendered using the ESPript server (3).

A

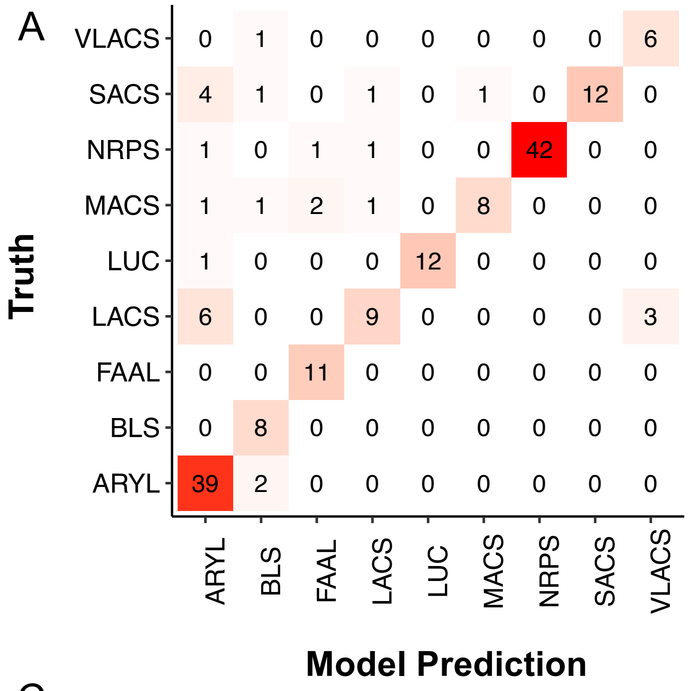


B

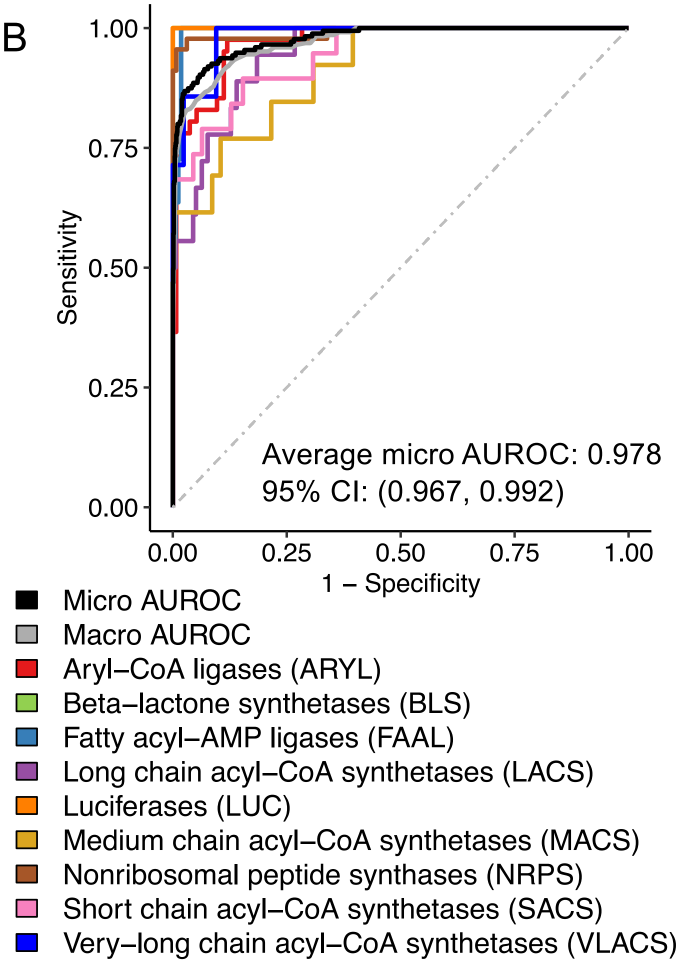


C

| **Test set classification accuracy** | |
|---|---|
| Random Forest | 0.833 ± 0.030 |
| Feeforward Neural Network | 0.827 ± 0.026 |
| Naïve Bayes | 0.820 ± 0.019 |

**Figure S2.** A) Confusion matrix of model predictions vs. truth for hold-out test set functional group classification using the random forest model B) Area under the receiver operating characteristic curve (AUROC) for enzyme function predictions. Colors correspond to different functions. C) Area under the receiver operating characteristics curve for substrate specificity predictions. Colors correspond to different functional classes as described in the legend as well as and macro (gray) and micro (black) AUROC averages.

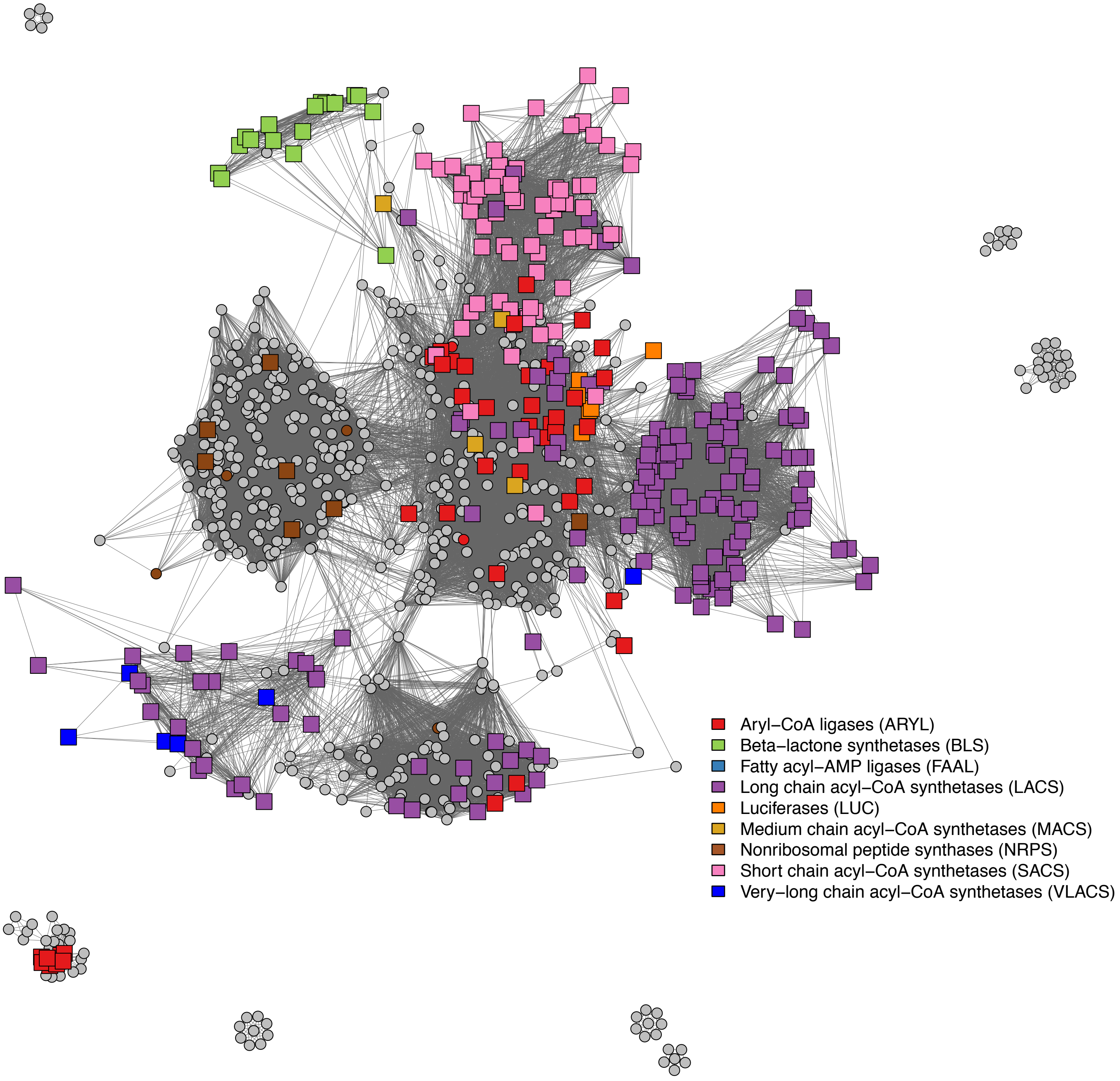


**Figure S3.** Sequence similarity network of MIBiG AMP-binding pHMM hits recovers a similar ‘hub’ topology to Fig. 3. Squares represent training set sequences and circles represent genes extracted from MIBiG biosynthetic clusters. The network was trimmed to a BLAST e-value threshold of 1 x $10^{-36}$.

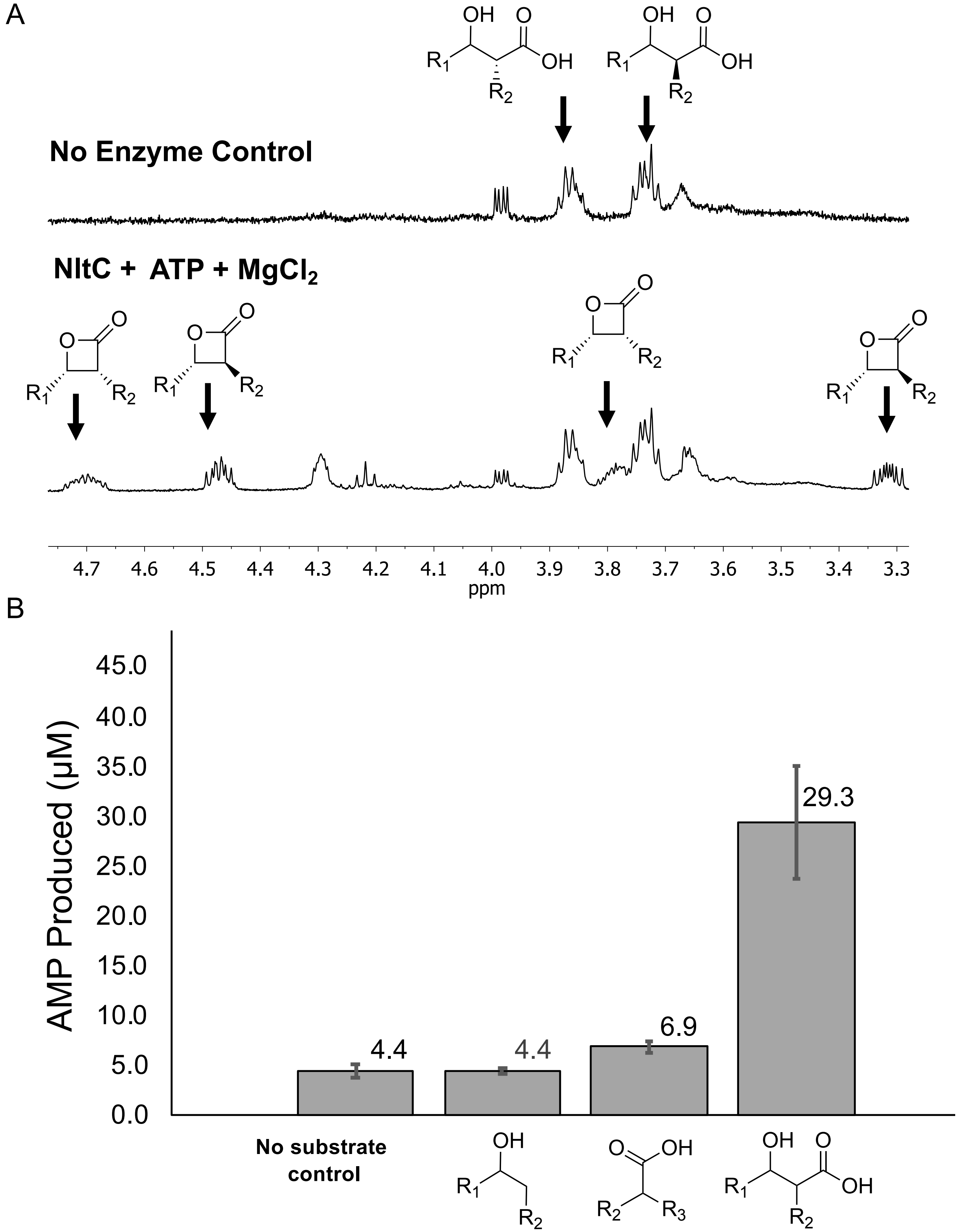


**Figure S4.** A) $^1$H-NMR of NltC-catalyzed formation of *cis*- and trans-β-lactones compared to no enzyme control. $R_1 = C_9H_{19}$, $R_2 = C_8H_{17}$. B) NltC activity with substrate mimics indicates that β-hydroxy acids are preferred. There is a slight stimulation of AMP release by the 2-hexyldecanoic acid substrate mimic compared to the no substrate control and secondary alcohol (10-nonadecanol). $R_1 = C_9H_{19}$, $R_2 = C_8H_{17}$, $R_3 = C_6H_7$.

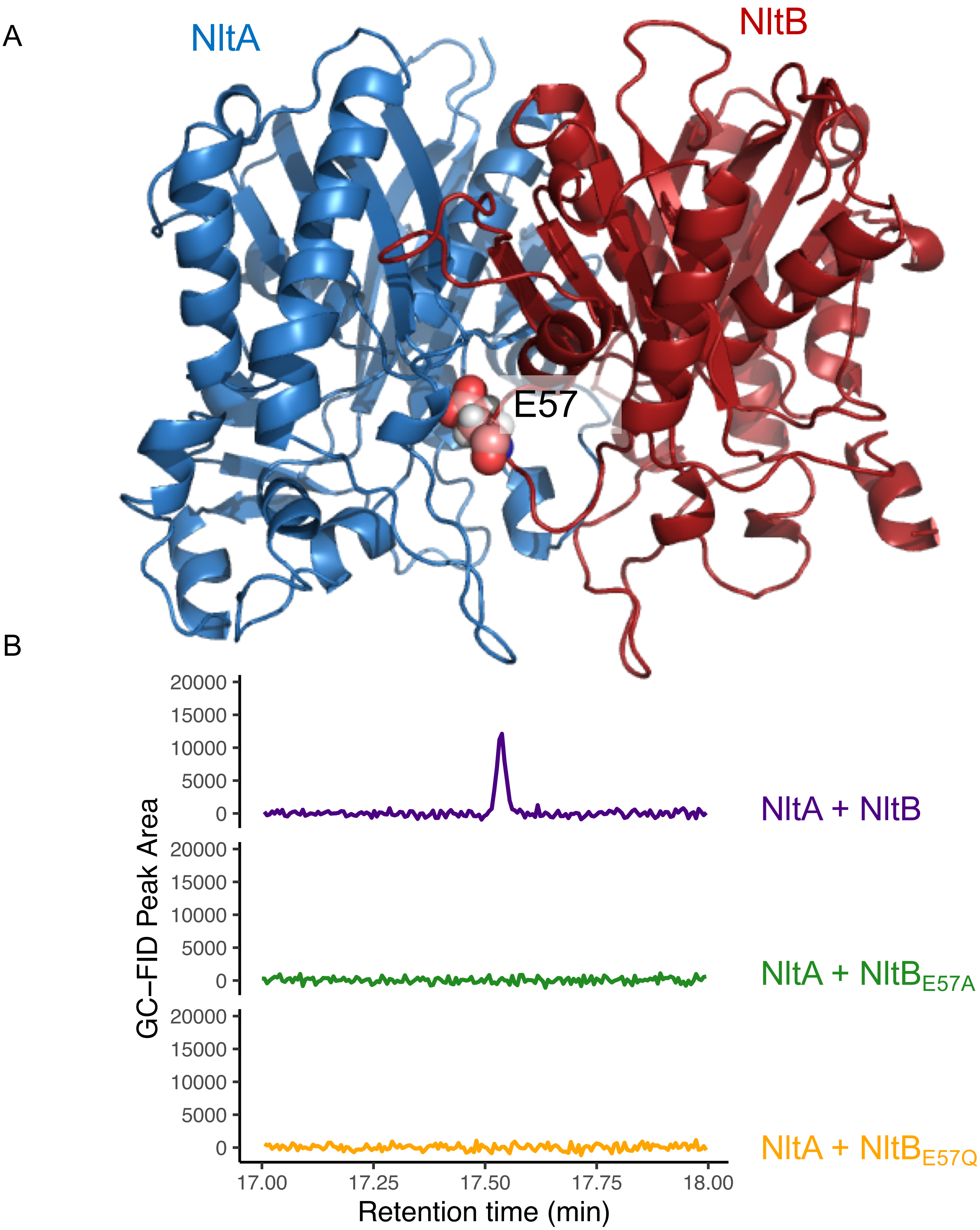


**Figure S5**. A) Homology model of NltA (red) and NltB (blue) heterodimer constructed using the Robetta server (4). B) NltA + NltB$_{E57A}$ (green) and NltA + NltB$_{E57Q}$ (orange) enzyme variants co-expressed do not condense decanoyl-CoA while NltA + NltB co-expressed (purple) are active.

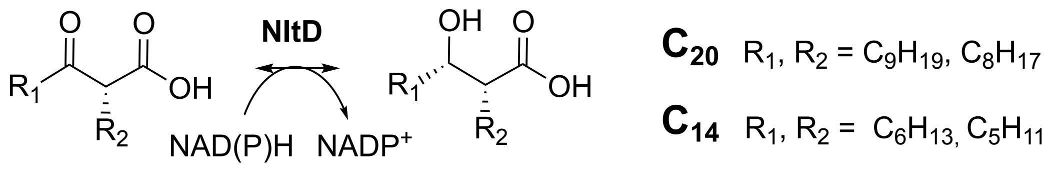


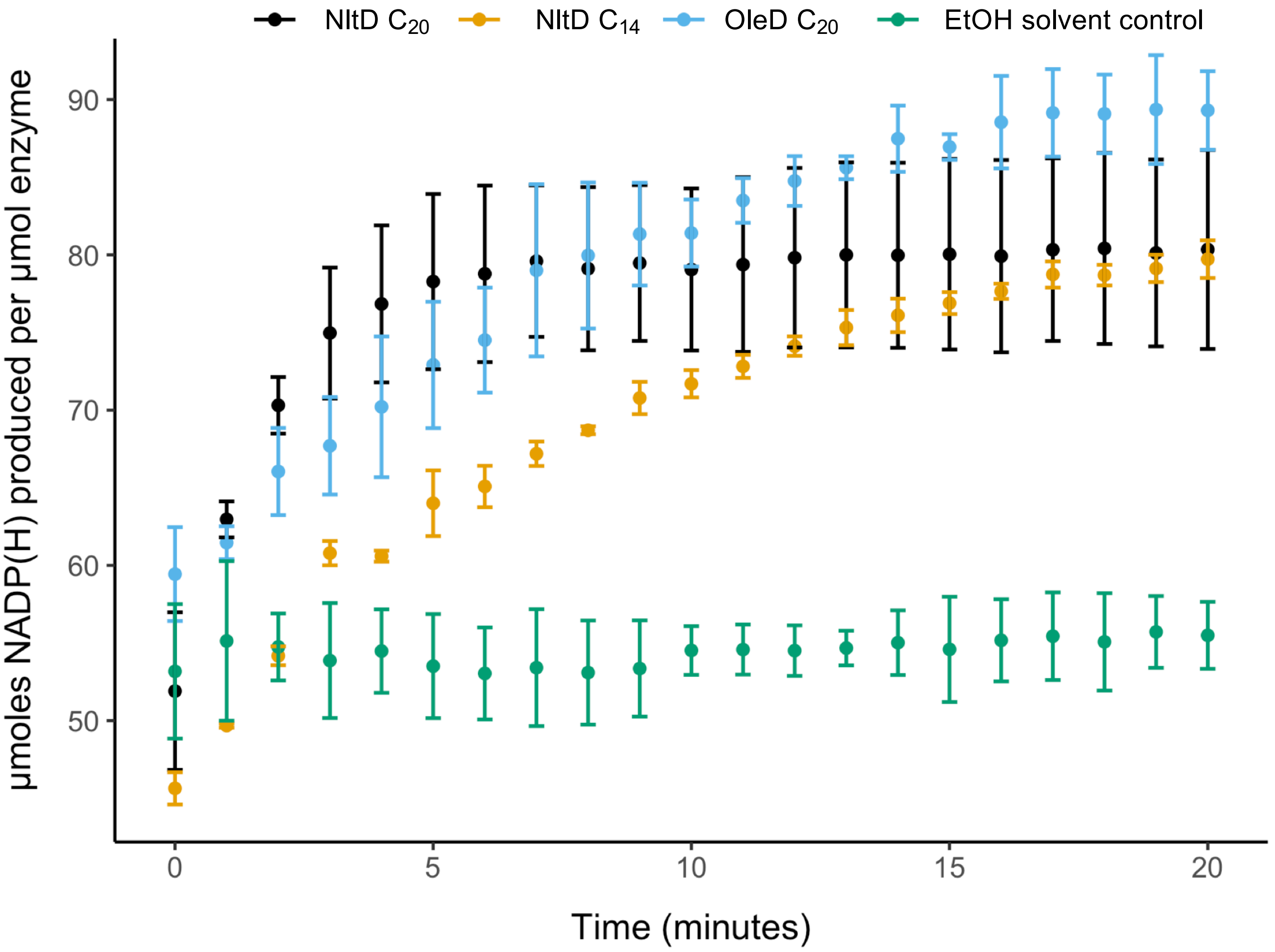


**Figure S6.** *Nocardia brasiliensis* NltD activity compared to a *Xanthomonas campestris* OleD (blue) positive control reveals NltD is active with both $C_{20}$ (black) and $C_{14}$ (orange) disubstituted β-hydroxy acids compared to a no substrate control with NltD (green).

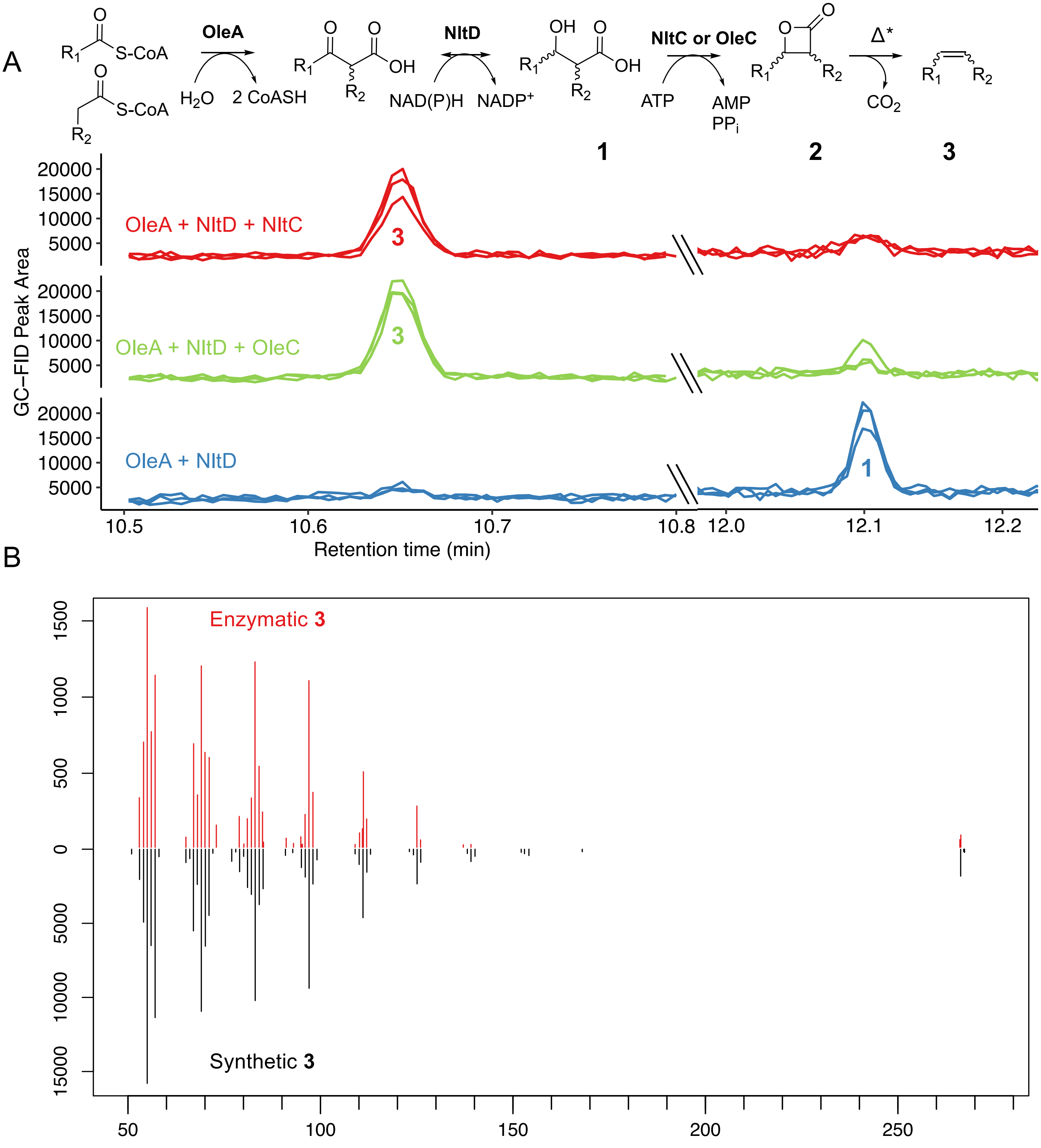


**Figure S7.** *In vitro* reconstitution of β-lactone biosynthesis pathway A) GC-MS traces indicate di-alkyl β-lactone products (**2**) are formed from a combination of three pathway enzymes (OleA, NltD, and NltC/OleC), ATP-$MgCl_2$, NAD(P)H, and decanoyl-CoA. $R_1$, $R_2$ = $C_9H_{19}$, $C_8H_{17}$. The addition of either NltC or OleC (β-lactone synthetases) are required to cyclize β-hydroxy acids (**1**) formed by OleA + NltD. B) Mirrored mass spectra of 9-nonadecene **(3)** resulting from thermal decarboxylation of synthetic **2** or enzymatically-formed **2**. n=3 for each enzyme combination. *In the heat of the GC inlet, β-lactones thermally decarboxylate to olefins. We used a program with an elevated inlet temperature for quantitative total conversion of β-lactones to olefins validated with a standard curve of synthetic β-lactones as described by Robinson et al. (5).

### Supporting file S1. Codon-optimized gene constructs for NltA and NltB

The following codon-optimized synthetic genes were cloned by Gibson assembly into standard pET28b+ vectors at NdeI/XhoI restriction sites and with N-terminal 6x-His tags.

Red = homology arms
Green = NdeI cut-site
Orange = XhoI cut-site

>WP_042260942.1 NltA [Nocardia brasiliensis]
tgccgcgcggcagccatATGAGAGCGTGGACATACGGCCCACGTCGCTTGCTGTTGCCGTCCGGCTCCGTCC
GTCGGACAGAGGATCAGCGTCGGAGTCGCAATCGCATATTCTCAACGATTTCGGGCGCGCGGGTCAG
CCACAGTCGCTTTGAGTCGATTGGGGCCTATCTGCCTTCAAACGTAGTCACAACTCAAGAGCTGTTAG
CGCGCCTGCAAGAGCCGCCCTCGTTTGATTTAGAGAAAATTACTGGGGTGAAGGAACGCAGAGTGCAC
GATACGAGACCTGAAAGCTACGAGGACAGCTTCGCTATTGCCATGAAGGCAGCACAGGATTGTTTGTC
GCGGAGTAGATACGAAGCTGCCGAGCTTGATGTAATAATAAGCACTTCAATAACCCGCAGTAACCACG
CAACTCGGATGTACATGGAACCTTCTTTCGCCGGTTCTATATCACGGCTTATAGGCGCCAAGAAGGCA
ATCACCTTCGATGTCAGTAACGCGTGTGCCGGGATGCTGACCGGAACCTATATTTTAGATCGTATGATA
CGGTCGGGTGCCGTGCGGAATGGTATGGTCGTGAGCGGAGAGGCTATCACGCCGATAGCTGACACTG
CCGTTGCGGAAATATCTGAAAAGTATGATTTGCAGTTTGCCAGTCTGAGTGTTGGTGATTCGGGGGCG
GCTGTTGTTTTAGACCAGGCTGTTGACGACGATGACAAGATACATTACATTGAGTTAATGACAGCATCT
GAGTATTCGCACCTTTGCTTGGGCATGCCTTCAGACAAGAGTCAAGGCGTTGCCTTGTATACAGATAAC
AGAAAAATGCATAATGAAGACCGGTTTTTGTTGGGCACCAACTCCCAGATCAACTATCTTGAAGCCCAG
GGCCGCACCTTTGCAGACGAGAAGTTCGACTACGTTATTCATCACCAATTTGGTGCGGCAGCTATTCC
GTGGATGAATGCAATCTGCGAGCGGGAGTTTGGCACGCCCATGCCGCCTGATTTGCGTGTAATAGAGA
AGTACGGGAACACGTCAACAACTTCTCATTTTATTGTGCTTCACGATCAGCTGAGCGAACAGAACATAC
CTGCGGGGTCGAAACTCTTAATGATTCCTGCCGCATCCGGCATCGTCACAGGTTTCTTATCGACGACG
ATCTCATCCCTTAAGGTATAActcgagcaccaccacca

>WP_042260944.1 NltB [Nocardia brasiliensis]
ctggtgccgcgcggcagccatATGGGGGTTCGCATCAAGTCTACCGGCGTGTCTCGCGCCGAAGATACGAATTC
CTCCGTGGAGAATTCAGGGCGCGCAGCTAAACAGTCATTGGAGCGCGCTGGGGTGCGCCCTGACCAG
GTCGGCGTCCTTATCAACGCAGGGGTATTCCGCGACTCTAATACGGTGGAACCAGCTGTATCTGCATT
GATCCAGAAGGCCGCCGGTATTGGTTTAGAATATGCAAAAGATGACCCGCGCACCTTCAGCTTCGATC
TTATGAATGGTGCAACTGGTGTCTTGAATGCGGTCGGGGTCGCAACAAGCATCCTTGAAACAGGATCG
GCGGAACATGTATTGATCGTCTCCGGTGATGCGCACCCCTCTATGACTCGCAAAGCAGCTACAGATGA
CTTTCCGTATGCCGCGTCGGGCGCCGCACTGCTTTTGGAACGTACTGACGAACCCGAGGGATTTGGA
CGCGTACATACCGTGAATGGCGAAGGGACGCCAGCGGTGGAGGCGTATGTCGACACGGCAACTATGG
GCTCGGTTGGCCGTGGTTTGATGACCGTAGAACGCGAACCAGATTTCGCCGCACGCTTGTTAAATGTA
GCGGTTGAGGCCGCACTGGCGGCCCTTAACGAGGCTGGACACGACGACCTTTCAGGGACCGCACTTA
TCGCATCTACACCCACGGCAGAATTTCCCTTACAGTTAGCAGACGCTCTTGGAATTGACGAGGATGCC
GTCCGTACCCCTGACTTAACCGATGGTGATCCTCACACGGCTGCTCTGCCACAAGCCTATCACCGCGC
CGTTGTTGATGAAACTTTACGCGAATTTGGACATGTGTTATTCGTGGCAGCTGGGGCAGGCCCCTCCG
CTGCCGCGGTGTTATATCGTCTGCCGGCATTAGCGGGAGCTACAGCCTAActcgagcaccaccaccaccaccact

>WP_067710538.1 NltA [Nocardia_yamanashiensis]
ccgcgcggcagccatATGCCCCATAGCCGCTTTGAGAGCATTGGGGCCTATCTGCCGACGAAGCGTGTGACC
ACTCAAGAACTGATAGCTCAGTTGAAAGAGGCTCCAGGTTTTGACCTGGAGCGTATTACAGGCGTAAA
AGAACGGCGGTTTAGAGACACAAGCCCAGATCACCATGAAGATAGTTTTACCCTGGCCATGAATGCGG
TGGAAGATTGTTTGTCTCGGAGCCAATACACTGCTGACGACATTGACGTGATTATATCCACCTCAATTA
CTCGCAGTAAAGACGGGACAAAAATGTACATGGAACCTAGCTTCGCTGGTAGTATCGCCAAAGCGATC
GGAGCAAGAGCGGACGTTATCACATTTGATTTGTCAAACGCCTGTGCGGGTATGCTTACAGGGACATA
TATATTAGACCGCATGATTAGAAGCGGTGCAGTACGTCGTGGTATTGTGGTTTCGGGCGAAGCCATCA
CTCCGATCGCAGAGACAGCGGTGGCCGAGATTTCCGAGAAATATGACCTTCAGTTCGCTTCGTTAACT
GTGGGGGACTCGGGTGCGGCCGTGGTACTGGATCAAGCCGTTGACGATAATGATAAGATACATTACAT
CGAACTGACCACAGCATCTGAGTATAGTCATTTATGTCTGGGTATGCCATCCGACAAATCACAAGGAGT
GGCTCTTTACACTGATAACCGCAAGATGCATAATGAGTCGAGATTTCTGCTGTGGACTGACACACAAGG
TGGCTATTTGGAGAGAGAGGGGCGCTCCTTTGCGGATGAGAATTTCGACTACATGATACACCACCAAT
TCGGAGCGGGCGCTATTCCTTTTATAAATGCAATCGCGGAACGTGAGTTTGGGGCTCCCATGCCGCCT
GACCTTAATGTCTTAGAGAAGTACGGGAATACTAGTACAACCTCACACTTCATTGTCCTTCATGATCAG
CTTAGCCAACAGGCTATTCCGACTGGTAGTAAACTGTTGATGGTGCCGGCAGCCAGTGGTGTTGTAGC
TGGATTCCTGTCCACCACGATTTCATCGTTAAAAGTGTAAgcttgcggccgcactcga

### Supporting file S2. NltC and NltD codon-optimized gene constructs

The following codon-optimized synthetic gene was cloned by Gibson assembly into a standard pET30b+ vector at NdeI/HindIII restriction sites and a C-terminal 6x-His tag.

Red = homology arms
Green = NdeI cut-site
Purple = HindIII cut-site

>WP_042260945.1 NltC *Nocardia brasiliensis*
gtttaactttaagaaggagatatacatATGTCTTCGGCCACGTATTGGCAAGCTATTGATCGTTTTCGTGCTTTTGC
CCGCGCAGAGCCGGACCGCGAGGCCGTAATTTATCCTGTTGGCACTGATGCCGCGGGCTTGCCAG
CTTATCGCCACATTTCTTATCGCGAGTTGGACGATTGGTCCGAAACAATCGCCGAACGTCTGACTGC
GAGCGGCGTCGGCAGTGGCACACGTACAATCGTCTTAGTCCTGCCTTCACCAGAACTTTATGCAAT
CTTATTCGCGTTATTGAAAATCGGGGCGGTCCCGGTAGTTATTGACCCAGGGATGGGACTTCGCAA
AATGGTTCACTGTTTACGCGCAGTTGAAGCAGAGGCATTCATCGGGATTCCACCAGCTCACGCAGT
ACGCGTCCTTTTTCGTCGCTCATTCCGCAAGGTTCGCACTACTGTAACAGTAGGAAAACGTTGGTTC
TGGCGTGGAGCAAAACTGGCAGCGTGGGGCACTACCCCGTCCGGGGGTGCGGTTGACCGCGTGC
CCGCGGACCCTGGTGATGTCTTGGTGATCGGCTTTACAACGGGCTCAACCGGACCAGCGAAAGCA
GTTGAATTAACTCACGGCAATTTAGCTTCTATGATTGATCAGGTACATACCGCACGTGGTGAGATTG
CACCCGAAACTAGTCTTATCACTTTACCGCTTGTTGGGATTCTTGACCTGTTGCTGGGATCTCGCTG
CGTCCTTCCCCCACTTATTCCCTCGAAGGTCGGGTCTACTGATCCTGCCCACGTTGCCCATGCGATT
GAAACGTTTGGTGTCCGCACCATGTTTGCATCACCAGCATTGCTTATTCCGCTGTTGCGTCATTTGG
AACAACAGCCGAATGAATTAAAAACGCTTGCGAGCATCTATTCGGGTGGGGCCCCGGTTCCAGATT
GGTGCATCGCAGGATTACGCGCGGCTTTAACGGATGATGTACAGATTTTTGCTGGCTACGGTAGTA
CTGAGGCATTACCGATGTCGCTGATCGAGAGCCGTGAACTTTTTGATGGACTGGTAGAACGTACTCA
CCGTGGGGAAGGCACGTGTATTGGCCGTCCCGCTGACCGCATCGATGCCCGTATCGTCGCTATCA
CGGATGACCCTATCCCGACCTGGGCGCGCGCAGAGGAGCTGGCCGGAGACCTTGCGCGCTCACG
CGGCATTGGTGAATTAGTCGTCGCTGGACCTAATGTATCGACTCACTACTACTGGCCTGATACGGCT
AACCGTCAGGGAAAGATCGTGGACGGTGACCGCATCTGGCATCGCACAGGAGATTTAGCGTGGATT
GACGACGCTGGGCGTATTTGGTTTTGTGGTCGTAAAAGCCAACGTGTAGTAACAGCTGACGGGCCT
ATGTTTACGGTACAAGTCGAACAAATCTTTAACACAGTCGCAGGAGTAGCGCGCACTGCGTTGGTAG
GGGTAGGGGCGCCGGGGGCACAGCGCCCCGTTTTGTGCATCGAGCTTAAACCAGATGCGGAGGG
AGCGGCGGTGGGCGCGGCGCTTCGTGCTCGTGGGGCTGAATTTGACTTAAGTCGCCCGATTGCTG
ATTTTCTGATTCATCCTGGCTTCCCGGTAGATATTCGCCACAATGCGAAGATCGGACGTGAGCAGCT
TGCACAGTGGGCGGGCGAACAGCTTGGCGCCCGTGCTaagcttgcggccgcactc

The following codon-optimized synthetic gene was cloned by Gibson assembly into a modified pMAL-c5x vector with a tobacco etch virus (TEV) protease cut site and a N-terminal 6x-His tag.

Blue = TEV protease cut site

>WP_042260949.1 NltD *Nocardia brasiliensis*
agggaaggatttcacatatggagaacctgtacttccaatcaATGTCGAAGGTCCTGGTGACCGGAGCAAGCGGTTTCCT
TGGAGGGGCGTTGGTACGCCGCTTGATCCGTGATGGAGCCCACGATGTGAGCATTTTGGTACGCC
GTACGTCTAATTTGGCTGATCTTGGCCCTGACGTAGACAAGGTTGAGCTGGTCTACGGCGATCTGA
CGGATGCTGCATCACTTGTTCAAGCTACATCCGGTGTCGACATTGTATTTCATTCGGCAGCACGCGT
TGACGAGCGTGGGACGCGTGAGCAATTCTGGCAAGAAAACGTACGTGCCACGGAATTGTTGCTGGA
CGCCGCGCGTCGCGGTGGTGCATCCGCCTTTGTGTTCATTTCGTCTCCCAGCGCACTGATGGATTA
TGACGGAGGCGACCAACTTGATATTGATGAGTCAGTACCGTATCCGCGCCGTTATCTGAATCTTTAT
TCCGAAACTAAGGCAGCCGCCGAGCGCGCGGTATTAGCCGCCGATACGACAGGATTCCGCACCTG
CGCACTGCGCCCGCGTGCTATTTGGGGGGCCGGCGACCGTTCCGGTCCCATTGTCCGTTTACTGG
GTCGCACAGGCACAGGCAAATTACCAGACATCAGCTTTGGACGTGACGTTTATGCATCTTTATGCCA
TGTGGACAATATTGTGGACGCCTGCGTGAAGGCCGCGGCTAACCCGGCGACAGTAGGAGGTAAAG
CCTATTTTATTGCAGATGCTGAAAAAACTAATGTGTGGGAGTTTCTGGGGGCGGTTGCGACTCGTCT
GGGCTACGAGCCTCCTTCACGTAAGCCCAATCCCAAGGTTATCGACGCCGTCGTCGGGGTAATTGA
GACAATTTGGCGTATCCCGGCAGTGGCGACACGTTGGTCCCCACCGCTTAGTCGTTACGCTGTAGC
CTTGATGACACGCTCCGCTACTTACGACACAGGGGCAGCAGCCCGTGATTTTGGATATCAACCAGT
GGTGGATCGTGAAACGGGGCTTGCGACCTTCTTGGCATGGTTGGAGAAGCAGGGGGGAGCAGTGG
AGTTAACCCGCACACTTCGCaagcttcaaataaaacgaaa
